# Na^+^/H^+^ antiporter activity by respiratory complex I controls mitochondrial Δψ and is impaired in LHON disease

**DOI:** 10.1101/2023.05.30.542932

**Authors:** Pablo Hernansanz-Agustín, Carmen Morales-Vidal, Enrique Calvo, Paolo Natale, Yolanda Martí-Mateos, Sara Natalia Jaroszewicz, José Luis Cabrera-Alarcón, Iván López-Montero, Jesús Vázquez, José Antonio Enríquez

## Abstract

The mitochondrial electron transport chain (mETC) converts the energy of substrate oxidation into a H^+^ electrochemical gradient (Δp), which is composed by an inner mitochondrial membrane (IMM) potential (ΔΨmt) and a pH gradient (ΔpH). So far, ΔΨmt has been assumed to be composed exclusively by H^+^. Mitochondrial Ca^2+^ and Na^+^ homeostasis, which are essential for cellular function, are controlled by exchangers and antiporters in the inner mitochondrial membrane (IMM). In the last few years, some of them have been identified, except for the Na^+^-specific mitochondrial Na^+^/H^+^ exchanger (mNHE). Here, using a rainbow of mitochondrial and nuclear genetic models, we have identified the P-module of complex I (CI) as the major mNHE. In turn, its activity creates a Na^+^ gradient across the IMM, parallel to ΔpH, which accounts for half of the ΔΨmt in coupled respiring mitochondria. We have also found that a deregulation of this mNHE function in CI, without affecting its enzymatic activity, occurs in Leber hereditary optic neuropathy (LHON), which has profound consequences in ΔΨmt and mitochondrial Ca^2+^ homeostasis and explains the previously unknown molecular pathogenesis of this neurodegenerative disease.

---

All organisms rely on the formation of transmembrane potentials to support energy balance, and the maintenance and regulation of these transmembrane potentials are crucial determinants of cell homeostasis. Eukaryotes rely on a plasma membrane potential and a mitochondrial inner membrane potential (ΔΨmt), the latter being particularly important for energy production and cell fate determination. The mETC is composed of several complexes and supercomplexes^1^. Mitochondrial complexes I (CI) and II (CII) respectively oxidize NADH and succinate to reduce ubiquinone (CoQ) to ubiquinol. Complex III (CIII) uses ubiquinol to reduce cytochrome c (cyt c), and complex IV (CIV) oxidizes cyt c to reduce O_2_ to H_2_O. These series of reactions are coupled to the translocation of H^+^ by CI, CIII, and CIV across the inner mitochondrial membrane (IMM) to form a H^+^-motive force (Δp). Δp, in turn, activates the phosphorylation of ADP to ATP, which is coupled to the electrophoretic entry of H^+^ through a fifth complex (CV). Δp is composed of an electrical component, ΔΨmt, which is negative in the mitochondrial matrix, and a chemical component, ΔpH, which is alkaline in the matrix and acidic in the intermembrane space^2^, with ΔΨmt accounting for approximately 80% of the total Δp^3^. CI is constituted by three structural modules^4-6^. The N-module mediates NADH oxidation and transfers the electrons to the Q-module, also called the CoQ-reducing module. The energy released by the NADH-CoQ oxidoreduction is transferred to the subunits in the H^+^ pumping module, or P-module. This is composed by mtDNA-encoded subunits which are evolutionarily related to the Na^+^/H^+^ antiporters of alkaliphilic and halophilic bacteria^7^.

Mitochondria also contain a panoply of exchangers and antiporters that allow them to maintain respiration, osmolarity, and volume, permit the entry and extrusion of substrates and metabolites and regulate allosterically many enzymes and transporters^8^. The mitochondrial Ca^2+^ uniporter (MCU) introduces Ca^2+^ into the mitochondrial matrix which, in turn, is extruded by the mitochondrial Na^+^/Ca^2+^ exchanger (NCLX) in an electrogenic exchange for 3 Na^+^ ions^9^. Intramitochondrial Na^+^ exit is mediated by a Na^+^-specific, highly active, electroneutral Na^+^/H^+^ exchanger (NHE), which extrudes Na^+^ and acidifies the mitochondrial matrix^10^. Notably, the molecular identity of the mitochondrial NHE remain to be identified^10^

## Respiratory complex I is the main mitochondrial Na^+^/H^+^ antiporter

In 2004, based on previous work in sequence similarity with alkalophilic bacterial Na^+^/H^+^ antiporters^7^, Stolpe and Friedrich reconstituted purified CI from *E*.*coli* into liposomes, and proposed that prokaryotic CI may be capable of a secondary Na^+^ antiport activity (i.e., This may occur in coordination with NADH:CoQ oxidoreduction and H^+^ pumping)^11^. Eight years later, Roberts and Hirst, using again purified CI from bovine heart and reconstituted in liposomes, proposed that also mitochondrial CI was capable of Na^+^/H^+^ exchange activity, but only after the enzyme transitioned to its deactive form (i.e., Na^+^/H^+^ antiport may not happen under the active, NADH:CoQ oxidoreduction-H^+^ pumping, state)^12^. These *in vitro* approaches suggested the possibility of CI being the molecular entity responsible for a mitochondrial NHE function. To evaluate this possibility, we first purified CI from pig heart mitochondria (Extended Data Figure 1a-d) and reconstituted it into proteoliposomes. In contrast to liposomes alone, CI proteoliposomes showed a strong Na^+^/H^+^ exchanger activity (Figure 1a and Extended Data Figure 1e-g).

**Figure 1.**
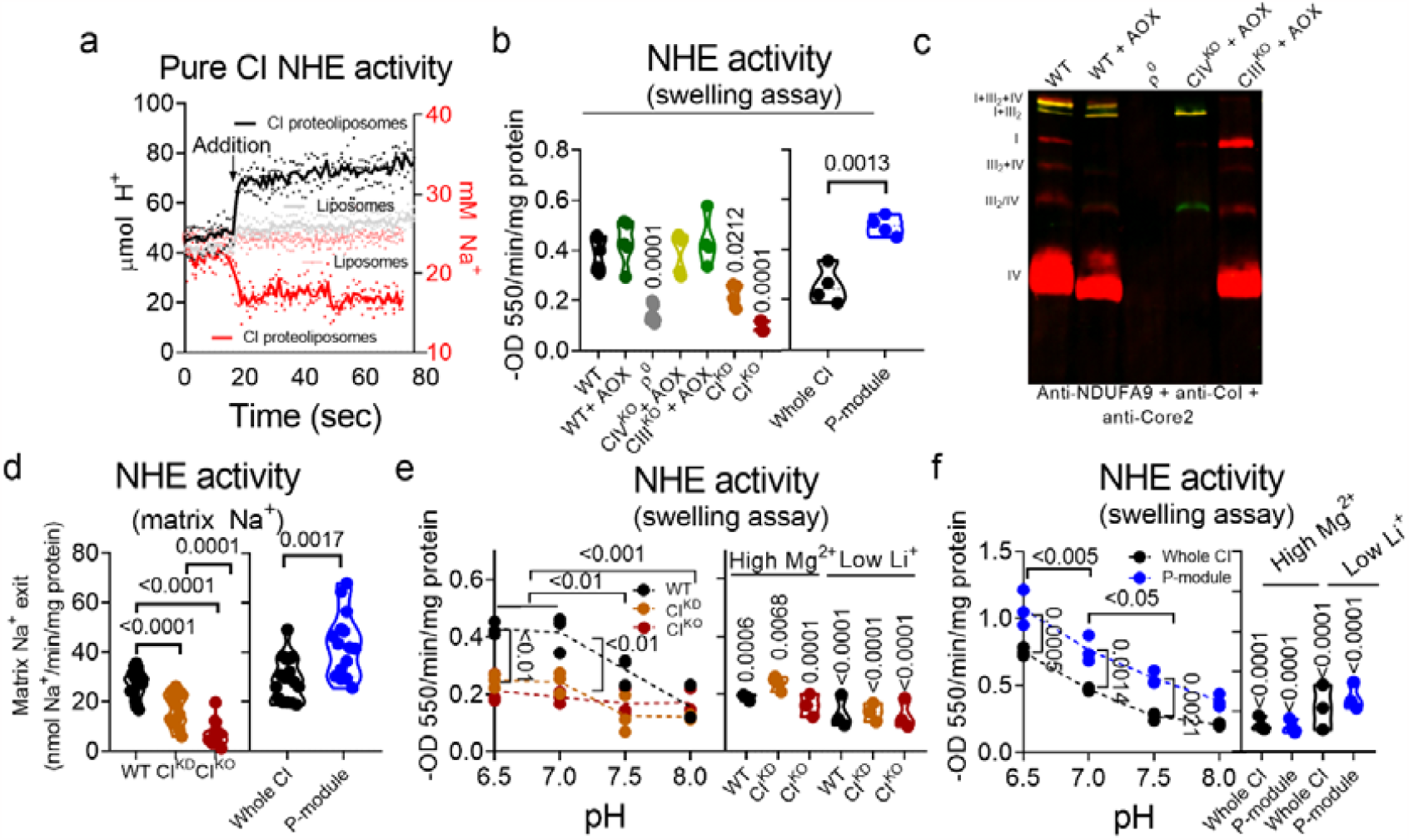
CI is the Na^+^-specific mitochondrial NHE. (**a**) (left axis) Buffer pH or (right axis) buffer Na^+^ content was measured before and after the addition of either liposomes (grey or pink) or pure CI-containing proteoliposomes (black or red; n=3). (**b**) Passive swelling NHE activity in WT, WT+AOX, ρ^0^ (CI, CIII and CIV absent), CIV^KO^+AOX (only CIV absent), CIII^KO^+AOX (only CIII absent), CI^KD^, CI^KO^ NDUFS4^WT^ and NDUFS4^KO^ non-respiring mitochondria (n=4). Only p-values against WT are shown. (**c**) BN-PAGE of WT, WT+AOX, ρ^0^ (CI, CIII and CIV absent), CIV^KO^+AOX (only CIV absent) and CIII^KO^+AOX (only CIII absent) mitochondria. Anti-NDUFA9 (CI) and anti-CoI (CIV) are colored in red, whereas anti-Core2 (CIII) is colored in green). (**d**) Passive NHE activity was measured as a readout of mitochondrial Na^+^ exit in WT, CICI^KD^, CI^KO^, NDUFS4^WT^ (Whole CI) and NDUFS4^KO^ (P-module; n=15; n=13 in CI^KO^). (**e**) Passive swelling NHE activity in WT, CICI^KD^ and CI^KO^ was assessed in different buffer pHs and in the presence of 50 mM MgCl_2_ or with 2 mM LiCl (n=3). (**f**) Passive swelling NHE activity in NDUFS4^WT^ (Whole CI) and NDUFS4^KO^ (P-module) was assessed in different buffer pHs and in the presence of 50 mM MgCl_2_ or with 2 mM LiCl (n=3). (d and e) p-values in “High Mg^2+^” and “Low Li^+^” are against WT or NDUFS4^WT^ (Whole CI) at their corresponding, same pH (i.e., pH 7).

Next, we wondered whether such CI-dependent activity could be measured in intact mitochondria. For that, we used several cellular knock-out (KO) models of different OXPHOS subunits: 1) a mutant with reduced levels of, but still present, CI or CI knock-down (CI^KD^)^13^, which only shows partial CI assembly by BN-PAGE (CI^KD^ in Extended Data Figure 2a); 2) a mutant with a complete loss of CI or CI^KO^, which show no detectable CI by BNGE (CI^KO^ in Extended Data Figure 2a)^14^; 3) and a KO model of the CI nuclear-encoded subunit NDUFS4 or NDUFS4^KO^, which has the assembly of the CI N-module severely impaired, leaving a CI subassembly with only the P module (named P-module; Extended Data Figure 2b); 4) a variety of cell lines lacking either all mtDNA-encoded OXPHOS complexes (ρº cells) or only CIII or CIV^15^.

CI^KD^, CI^KO^ and P-module cell lines displayed a significant decrease in CI activity (Extended Data Figure 2c and d). Notably, whereas CI^KD^ and CI^KO^ showed a reduction in mNHE proportional to their CI content, the only presence of the CI P-module produced a marked increase of mNHE (Figure 1b-c and Extended Data Figure 2e-j). Importantly, the reduction in mNHE activity could only be seen after the specific deletion of CI and not upon the removal of any other mETC complex (Figure 1b-d and Extended Data Figure 2a). All this indicates that intact mitochondria retain its mNHE function only when the P-module of CI is present, supporting the physiological implication of the NHE activity observed in isolated CI. In addition, these results also indicate that CI NADH:CoQ oxidoreductase function is not necessary for mNHE and that CI can exert this role in the absence of the N-module.

Classically, there have been two NHE types ascribed to mitochondria: 1) A highly active, Na^+^-specific NHE, inhibitable by low Li^+^, high Mg^2+^ and alkaline pH; 2) and a sluggish Na^+^-unspecific (i.e., it also catalyzes K^+^/H^+^ antiport) NHE, inhibitable by low Mg^2+^ and acidic pH^16-22^. We observed that neither of the CI deficient cell models was able to alter mitochondrial K^+^/H^+^ exchanger (KHE) function in intact mitochondria (Extended Data Figure 2k-l). Also, the pH-dependency profile showed that a rapid mNHE becomes inhibited with increasing buffer pH only in the presence of CI (Figure 1d). This pattern was still present in CI lacking its N-module (Figure 1e). Finally, CI P-module-dependent NHE activity was inhibited to similar values by high amounts of Mg^2+^ or low levels of Li^+^ (Figure 1e). Altogether, these results indicate that the P-module of mitochondrial CI constitutes the *bona-fide*, Na^+^-selective, mitochondrial Na^+^/H^+^ antiporter.

CI^KD^ and CI^KO^ cells represent a partial or total mNHE loss-of-function models, respectively, whereas the cells with only the CI P-module constitute a mNHE gain-of-function one, having all a similar decrease in NADH:CoQ enzymatic activity. Thus, we sought to characterize the bioenergetic impact of CI-mNHE activity in purified, intact mitochondria and whole cells. All cell models showed an increase in combined Antimycin A-dependent succinate:cyt c (i.e., CII+III) activity compared with their respective isogenic controls (Figure 2a and b and Extended Data Figure 3a, summarized in Extended Data Table 1). Increased CII+III activity is a well-known consequence of CI deficiency or inhibition^13^. CI^KO^ and mitochondria with only CI P-module, but not CI^KD^ ones, showed elevated CIV activity (Figure 2b and Extended Data Figure 3b), while CV activity, in forward or reverse mode, was higher only in mitochondria with CI P-module (Extended Data Figure 3c and d, summarized in Extended Data Table 1). Since CII is rate limiting for succinate oxidation^23^, it is expected that the increase in CII+III activity would be accompanied by a raise in succinate-dependent respiration and by an hyperpolarization of mitochondria. Surprisingly, only mitochondria with the P-module accomplished such expectation (Figure 2c-h and Extended Data Figure 3e-g, summarized in Extended Data Table 1). Intriguingly, mitochondria with only the CI P-module did not show lower respiration nor depolarization under CI substrates (Figure 2g and h and Extended Data Figure 3g), despite having lower CI activity (Extended Data Figure 2c and d). Although it is tempting to attribute these bioenergetic differences to the impact of the mutations in the NHE activity, alternative explanations need to be ruled out first.

**Figure 2.**
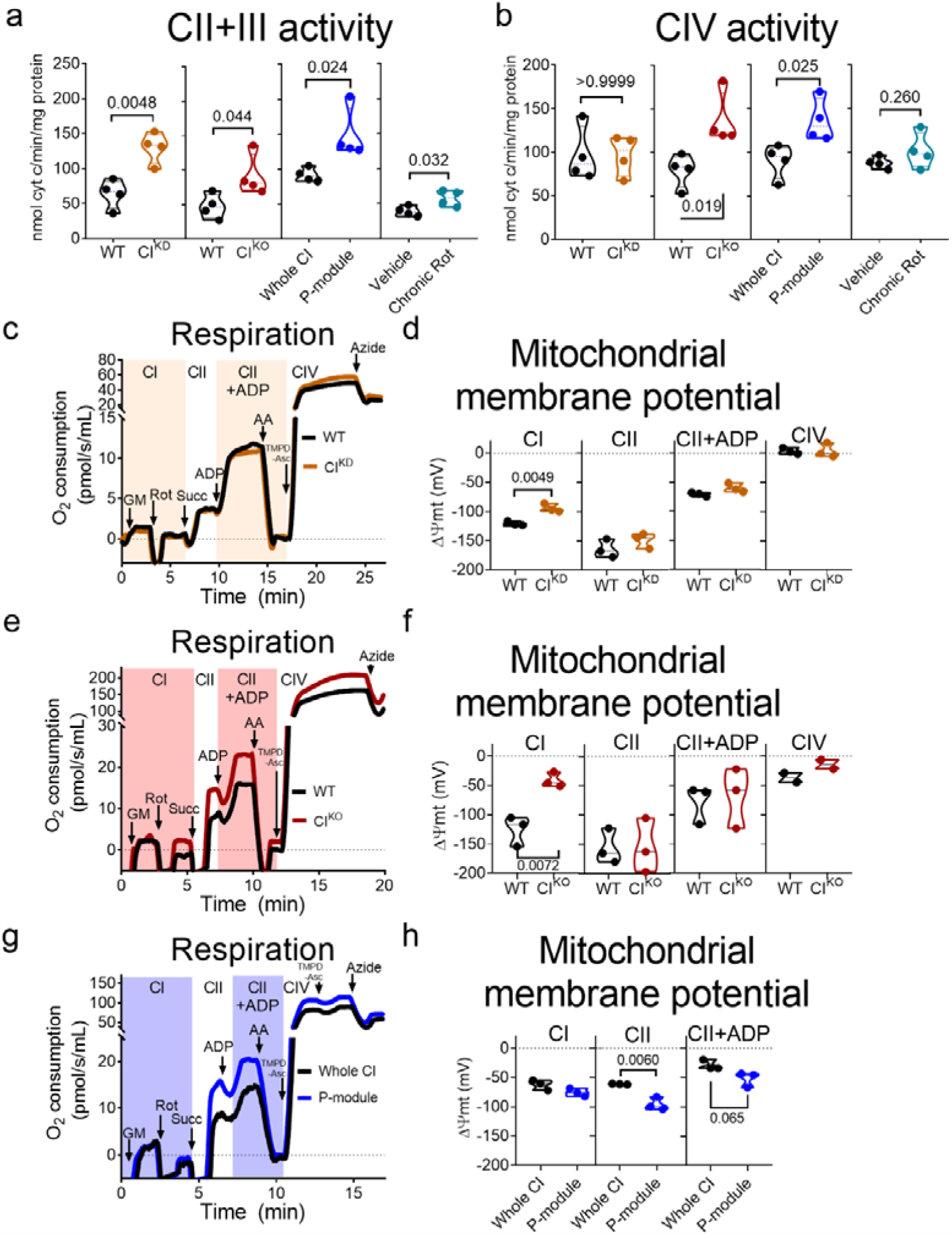
Characterization of CI-deficiency models. (**a**) Antimycin A (AA)-sensitive succinate-cyt c oxidoreductase activity in WT and CI^KD^, WT and CI^KO^, NDUFS4^WT^ (Whole CI) and NDUFS4^KO^ (P-module) and mitochondrial membranes from WT cells treated for 6 h with either vehicle or 250 nM rotenone (Chronic Rot; n=4). (**b**) Potassium cyanide (KCN)-sensitive cyt c oxidase activity in WT and CI^KD^, WT and CI^KO^, NDUFS4^WT^ (Whole CI) and NDUFS4^KO^ P-module) and Chronic Rot mitochondrial membranes (n=4). (**c, e** and **g**) Oxygen consumption rates in WT and CI^KD^ **c**, n=3), WT and CI^KO^ (**e**, n=3) and NDUFS4^WT^ (Whole CI) and NDUFS4^KO^ (P-module) isolated mitochondria (**g**, n=3). GM, glutamate/malate; Rot, 1 μM rotenone; Succ, succinate; ADP, adenosine di-phosphate; AA, 1 μM antimycin A; TMPD+Asc, N,N,N′,N′-tetramethyl-p-phenylenediamine + ascorbate; Azide, 50 mM, sodium azide. CI denotes CI-dependent respiration, CII denotes CII-dependent respiration, CII+ADP denotes CII+ADP-dependent respiration, and CIV denotes CIV-dependent respiration. (**d, f** and **h**) Calibrated TMRM signal in WT, CI^KD^ (**d**, n=3)), WT and CI^KO^ (**f**, =3; CIV n=2) and NDUFS4^WT^ (Whole CI) and NDUFS4^KO^ (P-module) cells (**h**, n=3), providing the CI+III+IV-ependent ΔΨmt (Panel CI), CII+III+IV-dependent ΔΨmt (Panel CII), CII+CIII+CIV+ADP-dependent ΔΨmt (Panel CII+ADP), and CIV-dependent ΔΨmt (Panel CIV).

The different CI deficient cell lines analyzed were obtained from two different mouse strains. CI^KD^ and CI^KO^ are cybrids with the nucleus of L929 cells, and NDUFS4^WT^ (labelled as CI^WT^) and NDUFS4^KO^ (labelled as CI^P-module^) are mouse adult fibroblasts isolated from C57BL/6J mice. We wondered whether this discrepant effect in their bioenergetic footprint may be due to differences in their origin or genetic background. To assess this, we chronically exposed cells with L929 nucleus and wild type mtDNA (WT) to a low dose of the CI inhibitor rotenone, leaving CI structurally intact. If the bioenergetic differences were due to the genetic background, we would expect that the increase in CII+CIII elicited by rotenone would not translate into a rise of succinate-dependent respiration and hyperpolarization.

We measured CII+III, CIV and CV activities (Figure 2a and b and Extended Data Figure 3c and d) and found that the elevation in CII+III promoted an increase in succinate-dependent respiration and hyperpolarization in mitochondria extracted from cells chronically exposed to a low dose of rotenone (Extended Data Figure 3h-i, summarized in Extended Data Table 1). Given that these results resemble that of those in mitochondria with only CI P-module, this points out that the observed bioenergetic differences are not due to their origin or genetic background. Next, we inquired if the fact that an increase in CII+III activity did not translate into a higher succinate-dependent hyperpolarization may be due to phenotypic differences between WT and CI^KD^ and CI^KO^ mitochondria in parameters known to affect bioenergetics, such as mitochondrial volume, ultrastructure or MIM permeability. We did not observe changes in mitochondrial volume (Extended Data Figure 4a-d), ultrastructure (Extended Data Figure 4c), fragmentation (Extended Data Figure 4d and e) or MIM permeability (Extended Data Figure 4f-i) which could explain the observed bioenergetic defects.

We also determined whether such differences between CI^KD^, CI^KO^ and mitochondria with only the P-module may be due to a succinate-specific effect and wondered whether they could be seen under different substrates. N,N,N,N-tetramethyl-p-phenylenediamine (TMPD)-Asc donate electrons to cyt c and provide a direct readout of respiration and ΔΨmt by CIV, avoiding possible effects of substate import to mitochondria and potential differences in electron flux through other, upstream complexes. Given their differences in isolated CIV activity (Figure 2b), we would expect hyperpolarization in CI^KO^ and mitochondria with only P-module, but not in CI^KD^. However, our expectations were again exclusively fulfilled by mitochondria with only P-module (Extended Data Figure 4j and k).

We wondered whether the observed effect may be because we were comparing mtDNA mutants (CI^KD^ and CI^KO^) against a knock-out of a nuclear encoded CI subunit (P-module). Thus, we studied the bioenergetic footprint and NHE activity comparing a different mtDNA mutant (ND4^KO^) and a knock-out of another nuclear encoded CI subunit (NDUFB11), both affecting CI P-module. All parameters measured in NDUFB11^KO^ and ND4^KO^ cells resembled those in CI^KD^ and CI^KO^: lower NHE activity (Extended Data Figure 5a), higher CII+III activity (Extended Data Figure 5b), similar degree of CI loss (Extended Data Figure 5c and d) and equal respiratory and ΔΨmt values than their isogenic counterparts (Extended Data Figure 5e-h). These results point out that the bioenergetic differences between the mitochondria with complete absence of CI and the ones with the only presence of the P-module are not explained by the fact that the CI subunit affected is encoded by the mtDNA or nuclear DNA or to a particular type of mutation.

To complete the characterization of the different cell models we performed proteomic studies of all cell lines. Principal component analysis of bulk proteomics revealed that the two different wild type cell lines grouped together despite their different genetic background (Extended data Figure 6a). In addition, wild type cells were well differentiated from all CI mutant cell lines (Extended data Figure 6a). Moreover, the CI^KD^ and CI^KO^ cells grouped very close, but clearly separated from the proteome of cells with only the CI P-module (Extended data Figure 6a). This was confirmed by gene set enrichment analysis (GSEA), which identified the gene ontology pathways significantly downregulated (Extended data Figure 6b) or upregulated (Extended data Figure 6c) in the CI deficient cells. In both cases these pathways were very similar between CI^KD^ and CI^KO^ and clear divergent from those of cells with only the CI P-module (Supplementary Table 1). Finally, proteomic analyses of isolated mitochondria confirmed that CI^KD^ and CI^KO^ had a progressive loss of all CI subunits, whereas mitochondria with only CI P-module retained the majority of its P-module subunits, as expected (Extended Data Figure 6d and h). No significant differences were observed in other mETC complexes (Extended Data Figure 6e-fg and i-k). Volcano plots show changes found in mitochondria from all cell models; however, we failed to see differences in the presence of the mitochondrial succinate carrier Slc25a10 or other proteins which may affect CII- and CIV-dependent respiration (Extended Data Figure 6l). In summary, no feature other than the different impact of NHE activity was associated with the divergent characteristic bioenergetic signature of the different CI deficient cell models.

## CI Na^+^/H^+^ antiport function regulates respiration and ΔΨmt

We noticed a correlation between CI-NHE activity and ΔΨmt. CI^KD^ and CI^KO^ displayed reduced CI-NHE activity and could neither increase the respiration nor hyperpolarize under CII or CIV respiration. On the other hand, mitochondria with only the P-module showed an elevated CI-NHE function and higher respiration and hyperpolarization in all cases (compare ΔΨmt and mNHE columns in Extended Data Table 1). We speculate that, as ΔΨmt controls respiration and this is exclusively composed by a H^+^ gradient, the observed differences could be a result of lower H^+^ ejected by CIII and/or CIV in CI^KD^ and CI^KO^. To assess this question, we measure H^+^ pumping directly in the same conditions as the previous experiments. As expected, H^+^ pumping was lower in all three models when they were reliant on CI substrates (CI-III-IV inset and traces in Extended Data Figure 7a and b). However, under CII substrates, CI^KD^ and CI^KO^ mitochondria showed higher H^+^ pumping than their isogenic control, resembling mitochondria with only the CI P-module (CII-III-IV inset, traces in Extended Data Figure 7a and b and summarized in Extended Data Table 1). This finding agrees well with the higher CII+III activity in all CI-deficiency models (compare Extended Data Figure 7a and b with Figure 2a) and further supports that mitochondrial succinate import is not differentially altered in the CI deficiency models. However, despite CI^KD^ and CI^KO^ mitochondria showed higher H^+^ pumping, this did not translate into a higher respiration or hyperpolarization. This intriguing contradiction may indicate that the Na^+^-selective CI-NHE function may be involved in the generation of ΔΨmt and control of respiration, so that ΔΨmt may be governed not only by a H^+^ gradient, but also by a Na^+^ gradient.

To evaluate this hypothesis, we first recorded ΔΨmt in WT and CI^KO^ mitochondria, supplied with CII substrates and upon the addition of the chemical Na^+^/H^+^ exchanger monensin to acutely restore a NHE function in isolated mitochondria. Whereas nigericin, a chemical K^+^/H^+^ exchanger, promoted hyperpolarization in isolated WT mitochondria (left inset in Extended Data Figure 7c), monensin did not (middle inset in Extended Data Figure 7c). In contrast, both drugs supported hyperpolarization of CI^KO^ mitochondria (left and middle insets in Extended Data Figure 7c), indicating that the restoration of a Na^+^ gradient by monensin addition contributed to the establishment of ΔΨmt in CI^KO^ mitochondria. No similar phenomenon was observed in mitochondria with only the CI P-module (right inset in Extended Data Figure 7c).

Second, we measured respiration rates and ΔΨmt in intact WT mitochondria maintained in an osmotically compensated, Na^+^-free buffer. The absence of Na^+^ slightly decreased the respiration rate and promoted mitochondrial depolarization by a third or a half, depending on the substrate conditions (Figure 3a and Extended Data Figure 7d). Mouse heart mitochondria behaved similarly (Extended Data Figure 7e and f). In parallel, the absence of Na^+^ promoted apparently higher CI- and CII-dependent H^+^ pumping in WT mitochondria, which we interpreted as the consequence of an inoperative NHE, unable to dissipate ΔpH (Figure 3b and c). None of these phenomena was observed in CI^KO^ mitochondria (Figure 3d-f and Extended Data Figure 7g), pointing to a role of the CI-NHE-dependent formation of ΔΨmt, possibly reliant on a Na^+^ gradient. To note, mitochondria respiring on CI substrates, in which CI is fully active, were able to use their CI-NHE function to contribute to the establishment of ΔΨmt (Figure 3a and Figure 3b), suggesting that fully active CI is still capable of performing a Na^+^/H^+^ antiporter activity. Indeed, we measured mNHE activity in mitochondrial samples with CI active or deactive and observed that, though deactive CI showed a clear increase in NHE, active CI was also able to perform it (Extended Data Figure 7h).

**Figure 3.**
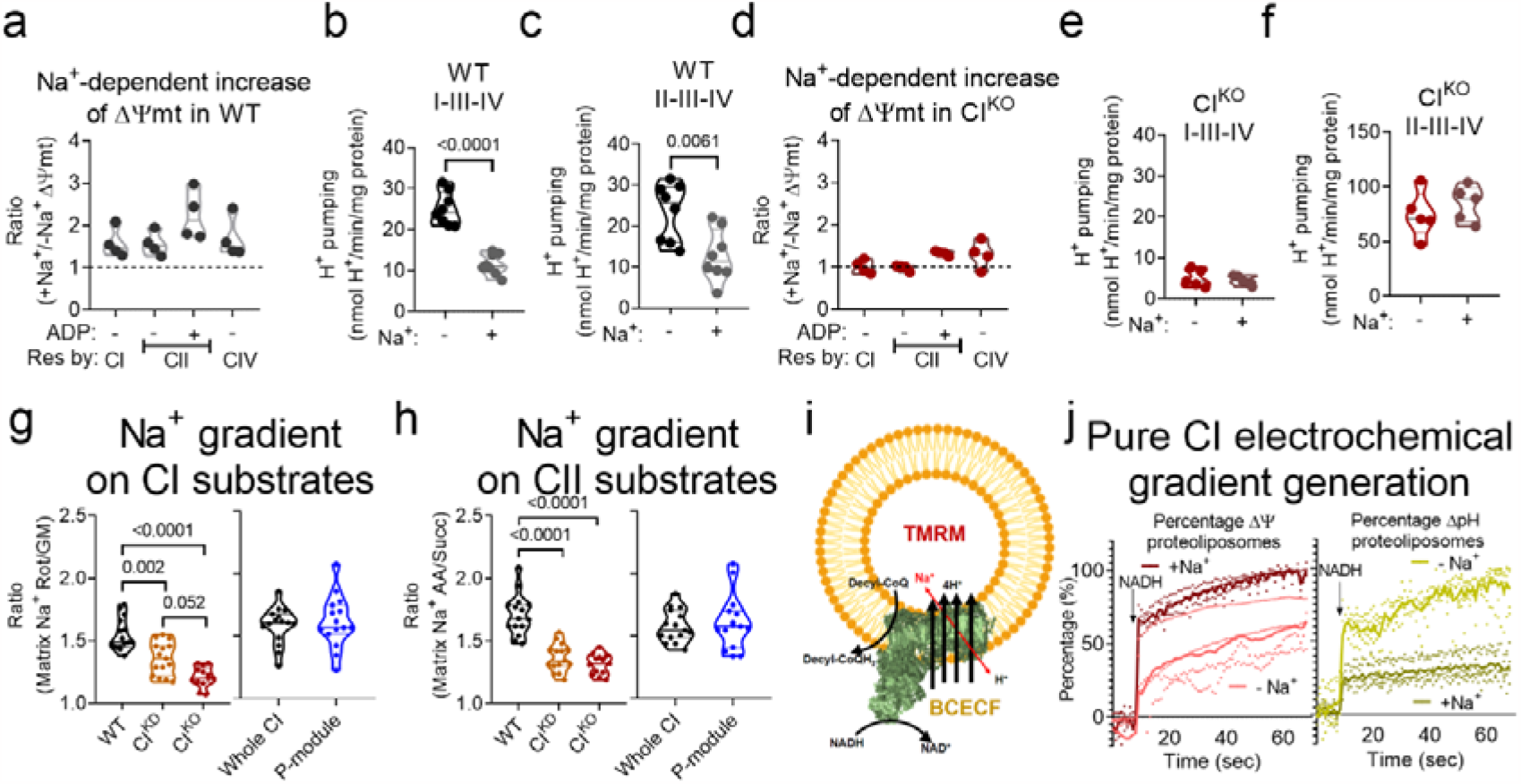
Mitochondrial Na^+^ gradient controls ΔΨmt. (**a**) Relative contribution of Na^+^ to ΔΨmt in WT mitochondria respiring on different substrates, calculated as the ratio of the calibrated TMRM signal in Na^+^-containing buffer to that in Na^+^-free buffer (n=4). (**b-c**) Mitochondrial matrix H^+^ pumping measured from the BCEFC-AM signal in WT isolated mitochondria respiring on CI (**b**) or CII (**c**) substrates in Na^+^-containing and Na^+^-free buffer (n=8). (**d**) Relative contribution of Na^+^ to ΔΨmt in CI^KO^ mitochondria respiring on CI, CII, CII+ADP, or CIV substrates, calculated as the ratio of the calibrated TMRM signal in Na^+^-containing to that in Na^+^-free buffer (n=4). (**e-f**) Mitochondrial matrix H^+^ pumping measured from the BCEFC-AM signal in CI^KO^ isolated mitochondria respiring on CI (**g**) or CII (**h**) substrates in Na^+^-containing and Na^+^-free buffer (n=5). (**g** and **h**) Na^+^ gradient in WT, CI^KD^, CI^KO^, NDUFS4^WT^ and NDUFS4^KO^ isolated mitochondria incubated with SBFI-AM and respiring on GM (**g**) or succinate (**h**), calculated as the ratio of the matrix Na^+^ concentrations determined after and before the addition of rotenone (**g**) or antimycin A (**h**) (n=15; n=13 in CI^KO^). (**i**) Scheme showing the experimental setup followed in section j. (**j**) Membrane potential (left inset; n=4) or intravesicular pH (right inset; n=5) of proteoliposomes reconstituted with pure CI were measured in a physiological buffer containing 130 µM decyl-CoQ, in the presence or absence of 100 mM NaCl and before and after the addition of 100 µM NADH.

We reasoned that if CI-NHE activity generates a Na^+^ gradient that contributes to the build-up of ΔΨmt and respiration, the presence of Na^+^ in the respiratory buffer would permit WT mitochondria respiratory and ΔΨmt values to equal those of CI^KO^ under CII or CIV substrates, despite the latter having larger H^+^ pumping. On the contrary, respiratory and ΔΨmt values should be lower in WT in the absence of Na^+^ since the Na^+^ gradient would not contribute to the build-up of ΔΨmt in these mitochondria (Extended Data Table 1). This is expected because CII-dependent H^+^ pumping in CI^KO^ mitochondria is higher than that of WT mitochondria. Indeed, CI^KO^ mitochondria showed higher respiration and hyperpolarization than WT in the absence of Na^+^ (Extended Data Figure 7i and j), which is in contrast with the very similar respiratory rates and ΔΨmt found in the presence of Na^+^ (Figure 2e-f and Extended Data Figure 3f). This result (i.e., CI^KO^ mitochondria having higher respiration and hyperpolarization) resembled the bioenergetic footprint of mitochondria with only the CI P-module and Chronic Rot mitochondria maintained in Na^+^-containing buffer (Extended Data Table 1).

Finally, the fact that we did not observe lower CI-dependent respiration and depolarization in mitochondria with only the P-module (CI inset in Figure 2g-e and Extended Data Figure 3g) may be due to the higher CI-NHE function in this model, as those experiments were carried out in the presence of Na^+^. Mitochondria with the CI P-module would build-up a Na^+^-dependent ΔΨmt sufficient to mask the lower CI activity when assessing O_2_ consumption and ΔΨmt. Thus, removing Na^+^ from the buffer may be enough to: 1) observe a lower CI respiration and depolarization mitochondria with only the CI P-module with respect to its WT counterpart; 2) depolarize mitochondria with only the P-module to slightly larger or even equal values than its isogenic WT, masking the effect of higher CII+III activity. Indeed, removing Na^+^ from the buffer revealed lower respiration and depolarization in mitochondria with only the CI P-module under CI substrates (Extended Data Figure 7k and l), making it consistent with its isolated CI activity. Similar results were obtained measuring H^+^ pumping (Extended Data Figure 7m). Furthermore, mitochondria with only the P-module oxidizing CII or CIV substrates were not able to raise respiration nor hyperpolarize mitochondria with respect to its isogenic WT in the absence of Na^+^ (Extended Data Figure 7k and l), despite having larger H^+^ pumping (Extended Data Figure 7m). This is because the contribution of the Na^+^ gradient to ΔΨmt in these mitochondria: 1) is larger than in its WT control; 2) and may be, in general, larger than that of the H^+^ gradient. To note, these results resemble those obtained with CI^KD^ and CI^KO^ in the presence of Na^+^ (i.e., mitochondria with only the CI P-module respiring without Na^+^ resemble CI^KD^/CI^KO^ with Na^+^), indicating that the presence or absence of Na^+^ in the respiratory buffer can mask many bioenergetically relevant features through its contribution to ΔΨmt.

In a third approach, we directly measured the mitochondrial Na^+^ gradient in isolated mitochondria. For respiration driven by CI or CII substrates, the Na^+^ gradient in CI^KD^ and CI^KO^ mitochondria was below-WT, proportionally to the level of assembled CI (Figure 3g and h). In contrast, the Na^+^ gradient in mitochondria with only the P-module was unaffected (Figure 3g and h), indicating that formation of the Na^+^ gradient requires a fully assembled CI P-module and CI-NHE function.

Fourth, if CI was able to build up a Na^+^ gradient and contribute to ΔΨmt due to its Na^+^-selective NHE function in intact mitochondria, NADH-oxidizing CI alone should reproduce such a behavior. To investigate this, pure CI from pig heart mitochondria reconstituted into liposomes was fed with NADH and decyl-CoQ to activate its H^+^ pumping function (Figure 3i) in the absence or presence of Na^+^. Active CI was able to create a membrane potential in the proteoliposomes through the increase in their H^+^ content in the absence of Na^+^ (pink lines of left inset and light green of the right inset in Figure 3j). In the presence of Na^+^, CI built up an even higher membrane potential at the expense of the H^+^ gradient generated by its own pumping function (red lines of left inset and dark green of the right inset in Figure 3j). These results corroborate that active CI contributes to membrane potential, not only by its H^+^ pumping function, but also through its NHE activity. In intact mitochondria, this H^+^- dissipating effect of CI-NHE increases respiration and contributes to the generation of ΔΨmt. This was absent in CI^KD^ and CI^KO^ models respiring under CII or CIV substrates, which is the reason why these models did not fulfill our expectations based on isolated CII+III/CIV activities. To corroborate that the absence of the H^+^ dissipating effect of CI-NHE conforms a brake for respiration, we measured respiration under different substrates in permeabilized mitochondria from CI^KD^ and CI^KO^. In these conditions, a higher CII- and CIV-dependent respiration could be readily seen (Extended Data Figure 7n and o), now confirming our expectations. Altogether, our results show that the CI-NHE function forms a Na^+^ gradient, at the expense of ΔpH dissipation, which raises respiration and contributes relevantly to ΔΨmt in mitochondria.

From the measurements taken throughout this study, we calculate that for respiration driven by CI or CII substrates the Na^+^ gradient contributes around one third of the total ΔΨmt (Extended Data Table 2). However, when CV is activated by the presence of ADP (state 3) and partially dissipates ΔpH, the electrical contribution of the Na^+^ gradient reaches approximately half of the ΔΨmt (Extended Data Table 2). The mitochondrial Na^+^ gradient thus either parallels the mitochondrial H^+^ gradient (compare WT in Extended Data Figure 8a and b with WT in Figure 3g and h) or even surpasses it (compare P-module in Extended Data Figure 8c and d with Whole CI in Figure 3g and h). Such a contribution of the Na^+^ gradient to ΔΨmt in coupled-respiring mitochondria, together with its effects in respiration, were also obvious in intact cells (Extended Data Figure 8e-i).

## The complex I Na^+^/H^+^ antiport function is specifically damaged in the mtDNA 11778G>A LHON mutation

The fact that, the two function of CI, NADH:CoQ oxidoreductase and mNHE are independent, raised the possibility that mutations in key CI P-module residues, altering CI NHE activity may alter ΔΨmt, independently of their CI NADH:CoQ oxidoreductase activity and H^+^ pumping capacity. LHON is a mitochondrial disorder causing central vision loss at early age by degeneration of the optic nerve and it is produced by mutations in mitochondrial DNA (mtDNA)-encoded CI subunits. Particularly, m.11778G>A (Supplementary Video 1), the most common mutation producing LHON, do not show a detectable decrease in neither CI activity^24^,^25^ (Figure 4a-b), CI-III-IV-dependent H^+^ pumping (Figure 4c) nor CI assembly (Extended Data Figure 9a). Likewise happened to CI^KD^, CI^KO^ and cells with only the CI P-module, LHON mitochondria displayed higher CII+III activity (Figure 4d) and, similarly to CI^KD^, comparable CIV activity than its control (Extended Data Figure 9b), which turned into larger CII-III-IV-dependent H^+^ pumping (Figure 4e). However, LHON mitochondria showed lower Na^+^-specific mNHE activity compared to its isogenic control (Figure 4f and Extended Data Figure 9c-e), which translated into a lower Na^+^ gradient (Figure 4g and h), mitochondrial depolarization (Figure 4i) and decreased respiration under CI substrates (Figure 4j). These results point out that a specific mutation in a mtDNA-encoded CI subunit, which do not interfere with its NADH:CoQ oxidoreductase activity, its assembly nor its H^+^ pumping capacity, and only affects its CI-NHE function, is able to promote defects in the maintenance of ΔΨmt andrespiration, also in intact cells (Extended Data Figure 9f and g).

**Figure 4.**
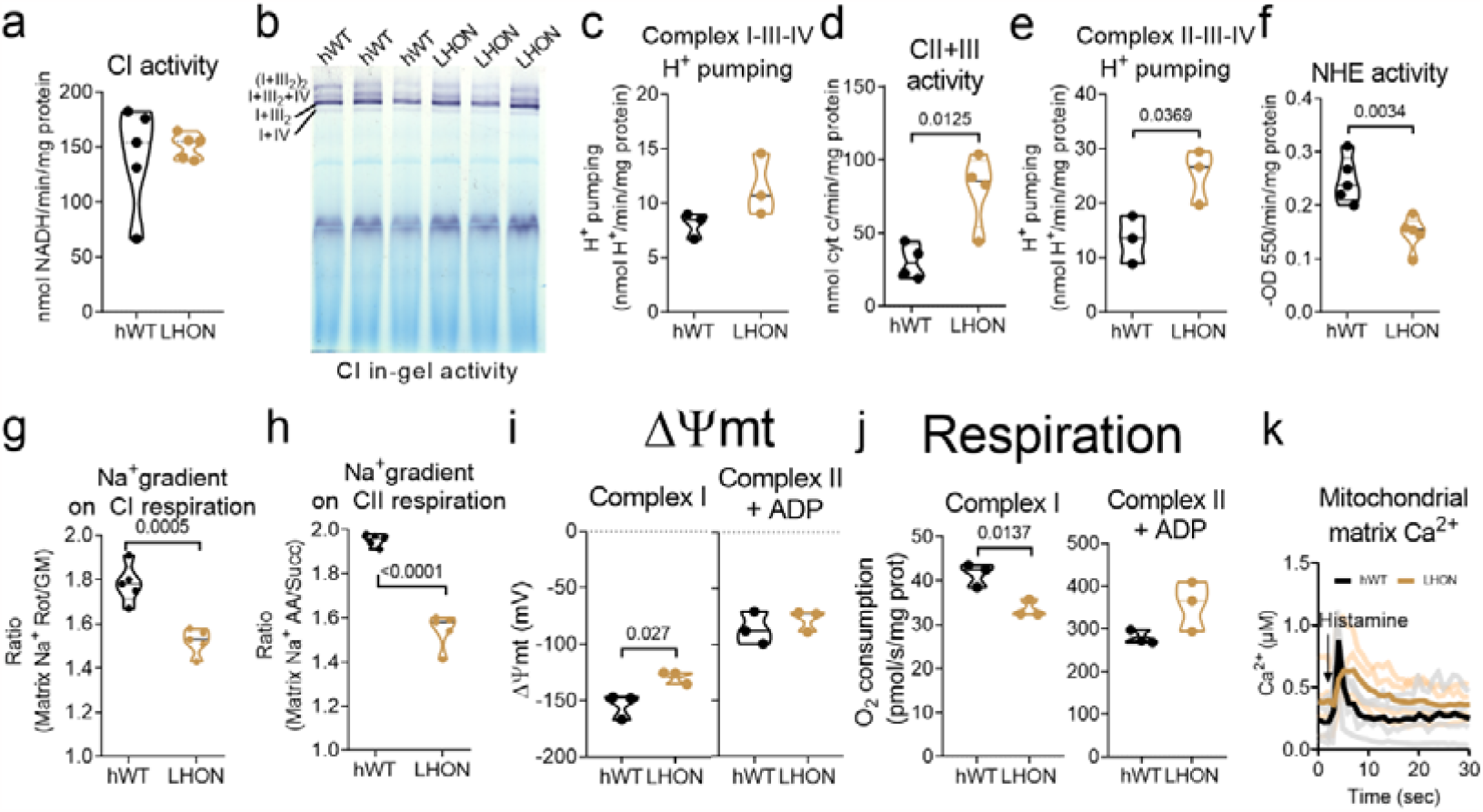
CI-NHE activity is specifically altered in LHON. (**a**) Rotenone-sensitive NADH-decylCoQ oxidoreductase activity in LHON vs control mitochondrial membranes (n=5). (**b**) CI in-gel activity in control (hWT) and LHON mitochondrial samples. (**c**) Mitochondrial matrix H^+^ pumping measured from the BCEFC-AM signal in hWT and LHON isolated mitochondria respiring on CI substrates in Na^+^-containing buffer (n=3). (**d**) Antimycin A-sensitive succinate-cyt c oxidoreductase activity in LHON vs hWT mitochondrial membranes (n=4). (**e**) Mitochondrial matrix H^+^ pumping measured from the BCEFC-AM signal in hWT and LHON isolated mitochondria respiring on CII substrates in Na^+^-containing buffer (n=3). (**f**) Passive swelling NHE activity measured in hWT vs LHON mitochondria (n=5). (**g-h**) Na^+^ gradient in hWT versus LHON isolated mitochondria, incubated with SBFI-AM and respiring on GM (**g**) or succinate (**h**) and calculated as the ratio of the matrix Na^+^ concentrations determined after and before the addition of rotenone (**g**) or antimycin A (**h**; n=5). (**i**-**j**) Calibrated TMRM signal (**i**) and respiration (**j**) in hWT versus LHON respiring with CI or CII+ADP substrates (n=3). (**k**) Mitochondrial Ca^2+^ was measured with calibrated Cepia2mt in hWT vs LHON cells before and after stimulation with 100 µM histamine (n=4). respiration, also in intact cells (Extended Data Figure 9f and g).

An important factor for the pathogenesis of LHON is the deregulation of mitochondrial Ca^2+^ management, which plays a particularly important role in the opening of the mitochondrial permeability transition pore (mPTP)^26^. Given that mitochondrial Ca^2+^ homeostasis is tightly linked to mitochondrial Na^+^ homeostasis through the roles of mNHE and NCLX, we wondered whether such a defect in CI-NHE could lie behind the known defects of LHON mitochondria to handle Ca^2+^. As such, given that mNHE function is impaired in LHON, mitochondrial Ca^2+^ entry would be expected to induce a slighter pH decrease in the matrix than its WT counterpart. Basal mitochondrial Ca^2+^ levels were higher in LHON (Extended Data Figure 9h). Also, both mitochondrial Ca^2+^ entry and exit were diminished (Figure 4k and Extended Data Figure 9i and j), which were not due to alterations in the levels of proteins involved in the homeostasis of mitochondrial Ca^2+^ (Extended Data Figure 9k and l) or lower cytosolic Ca^2+^ entry (Extended Data Figure 9m). We measured mitochondrial matrix pH and observed that, basally, LHON matrix was more acidic (Extended Data Figure 9n), presumably due to higher CV content and activity (Extended Data Figure 9a and n), and that the pH decrease during mitochondrial Ca^2+^ entry, which is caused by the activity of mNHE coupling Na^+^ exit with H^+^ entry, was also slighter (Extended Data Figure 9n). In fact, a similar, or even a deeper phenotype could be observed in CI^KO^ (Extended Data Figure 9p and q). All these results show that the specific decrease in CI-NHE activity in LHON not only directly affects ΔΨmt and respiration by depolarizing mitochondria and lowering oxygen consumption, but also alters mitochondrial Ca^2+^ management, causally explaining the molecular pathogenesis of a disease-causing mutation with a previously unknown etiology. Indeed, artificial restoration of Na^+^/H^+^ antiport with monensin was sufficient to reverse the bioenergetic phenotype in LHON mitochondria (Extended Data Figure 9r).

All in all, we provide here strong evidence supporting that CI conducts the mitochondrial Na^+^-specific mNHE function under both active and deactive forms. The presence of CI-NHE activity as well as the concomitant formation of a mitochondrial Na^+^ gradient enables the control of ΔΨmt, in addition to ΔpH, and a more efficient use of substrates in terms of oxygen consumption. Indeed, a specific deficiency in the CI-NHE function, independently of CI NADH:CoQ oxidoreductase activity, contributes to the molecular mechanism of the m.11778G>A mutation at CI through deregulation of mitochondrial Ca^2+^ homeostasis, which is responsible for the mtDNA-linked LHON.The discovery that the Na^+^ gradient controls ΔΨmt, together with its tight regulation by a non-canonical NHE function of CI, introduces a new and unexpected layer of regulation to mitochondrial bioenergetics, with extensive implications for neurological physiology and disease.

## Acknowledgments

We thank Dr. Sara Cogliati (CMBSO, UAM-CSIC) for helping with TEM imaging; Andrea Curtabbi, Dr. Demetrio Julián Santiago Castillo, Dr. Silvia Priori (CNIC), Dr. Eduardo Rial (CIB-CSIC) for fruitful discussion, and M. M. Muñoz-Hernandez, R. Martínez de Mena, E.R. Martínez Jiménez, and C. Jiménez for technical assistance. Scheme figures were made with BioRender. PN and ILM would like to thank Dr. Luis Calvo Adiego from the Icarlopsa Slaughterhouse for the generous gift of fresh porcine heart.

## Author contributions

Conceptualization: PHA and JAE. Methodology: PHA, CM (BN-PAGE of pure CI and OXPHOS mutants), YMM (Opa1 western blot), PN, ILM (Purification and reconstitution of CI in proteoliposomes) and EC (Proteomics). Investigation: PHA. Funding acquisition: JAE and JV. Project administration: JAE. Writing: PHA and JAE. Writing – review & editing: PHA, YMM, SJ, PN, ILM, EC, JV and JAE. PN and ILM would like to thank Dr. Luis Calvo Adiego from the Icarlopsa Slaughterhouse for the generous gift of fresh porcine heart. Microscopy experiments were performed in the Microscopy and Dynamic Imaging Unit.

## Competing interests’ declaration

Authors declare that they have no competing interests.

## Additional information

## Funding

This study was supported by competitive grants from the *Ministerio de Ciencia e Innovación* (MCIN) RTI2018-099357-B-I00, and CIBERFES (CB16/10/00282), Human Frontier Science Program (grant RGP0016/2018), and Leducq Transatlantic Networks (17CVD04) to JAE. PHA is supported by a JdC IJC2020-042679-I, YMM is supported by a FPI-SO fellowship PRE2018-083478. The CNIC has the support of the Carlos III Health Institute (ISCIII), the Ministry of Science and Innovation (MCIN), the Pro CNIC Foundation and is a Severo Ochoa Center of Excellence (CEX2020-001041-S financed by MICIN/AEI / 10.13039/501100011033). ILM acknowledges financial support through the TECNOLOGíAS 2018 program funded by the Regional Government of Madrid (Grant S2018/BAA-4403 SINOXPHOS-CM).

## Data and materials availability

All data are available in the main text or the supplementary materials. Cell lines will be available under a materials transfer agreement (MTAs).

## Notes

### Competing Interest Statement

The authors have declared no competing interest.

### Summary of Updates

The present version of the manuscript has been revised to add several significant results: - We characterize the type of Na/H antiporter that CI is. We have found that CI is the Na-specific mitochondrial Na/H antiporter. - Reconstitution of pure CI into proteoliposomes which were able to perform Na/H antiporter activity. When fed with CI substrates, such proteoliposomes were able to maintain a Na-dependent membrane potential. - Results regarding Ca management in LHON have been provided. We show causality between impairment of mitochondrial Na/H activity and Ca homeostasis in LHON and CI deficiency model.

