## Supplementary material for "Na^+^/H^+^ antiporter activity by respiratory complex I controls mitochondrial Δψ and is impaired in LHON disease"

### Materials and Methods

#### Cell culture and treatment

Mouse WT, $CI^{KD}$, $CI^{KO}$ and $ND4^{KO}$ cybrids were generated in the lab as described elsewhere. $CI^{KD}$ carries a frameshift mutation (13887iC) that created a stop codon 51 to 53 bp downstream of a 6-C stretch within the *mt-ND6* gene, resulting in a 79-amino-acid truncated polypeptide. $CI^{KO}$ consisted of the deletion of an A in a track of 7 As at position 10227 in the *mt-ND6* gene. This mutation led to a truncated polypeptide of only 26 amino acids (PMID: 20385768). WT were generated with untreated cells and following the same protocol for cybrid generation. hWT and LHON cybrids were generated by following the protocol for transmitochondrial cybrid generation on 143B cells and mitochondria from either healthy individuals or healthy patients. $NDUFS4^{WT}$ and $NDUFS4^{KO}$ were generated from $NDUFS4^{WT}$ or $NDUFS4^{KO}$ adult mouse (kindly donated by Dr. David Sancho), respectively. $NDUFB11^{WT}$ and $NDUFB11^{KO}$ were generated from mouse embryo and CRISPR-Cas9 technology (kindly donated by Dr. Alexandra Trifunovic).

WT, $CI^{KD}$, $CI^{KO}$, $ND4^{KO}$, hWT and LHON cybrid cells, and $NDUFS4^{WT}$, $NDUFS4^{KO}$, $NDUFB11^{WT}$ and $NDUFB11^{KO}$ mouse embryonic fibroblasts (MEFs), were cultured in DMEM supplemented with 5% FBS, 100 U/mL penicillin, and 100 μg/mL streptomycin. All cells were correctly identified by proteomic analysis, CI assembly assessed by Blue Native polyacrylamide gel electrophoresis (BN-PAGE), and CI spectrophotometric activity. Chronic rotenone (Chronic Rot) treatment consisted of incubating WT cybrids with 250 nM rotenone for 4-6 hours. This rotenone concentration was maintained throughout the detachment procedure in all buffers (trypsin and PBS) until the cell pellet was placed on ice, since removing rotenone or increasing its content during detachment was found to uncouple mitochondria. All cultures were routinely checked for mycoplasma contamination and tested negative.

#### Mitochondria isolation

Mitochondria were isolated from cybrids, MAFs, or mouse heart using a protocol adapted for either cell culture or mouse tissues[26]. Briefly, hearts and liver were cleaned of blood by rinsing in ice-cold phosphate buffered solution (PBS), and the tissue was disrupted with a blade on ice. Disrupted tissue, cybrids, or MAFs were resuspended in a buffer containing 0.32 M sucrose, 10 mM Tris-base and 1 mM EDTA (pH 7.4), placed in a glass Elvehjem potter, and homogenized by up and down strokes using a motor-driven Teflon pestle. Successive homogenization–centrifugation steps yielded the mitochondria-containing fraction, which was quantified with the Bradford protein assay.

#### Mitochondrial membrane isolation and measurement of complex activities

Mitochondrial membranes from cultured cells or liver were obtained after freeze–thawing isolated mitochondria, and OXPHOS enzyme activity was measured as previously described[27], using between 0.5 and 20 μg protein per sample, depending on the activity to be measured. Rotenone-sensitive NADH-ubiquinone decylubiquinone (decyl-CoQ) oxidoreduction (CI activity) was measured by changes in absorbance at 340 nm. Antimycin A-sensitive succinate-cyt c oxidoreductase activity (CII + CIII activity) was calculated after measuring changes in absorbance at 550 nm. Potassium cyanide (KCN)-sensitive cyt c-oxygen oxidoreductase activity (CIV activity) was calculated after measuring changes in absorbance at 550 nm. Oligomycin-sensitive ATPase activity (CV activity) was calculated after measuring changes in absorbance at 340 nm driven by the pyruvate kinase reaction coupled to ADP phosphorylation by CV.

##### Measurement of $O_2$ consumption and mitochondrial membrane potential

Measurements were made with an O2k Oxygraph instrument (Oroboros Instruments) with the attached fluorescent module. Tetramethylrhodamine methyl ester (TMRM) in quenching mode (2 µM) was used for the detection of mitochondrial membrane potential. Oxygen and TMRM calibration were performed before every experiment. Cells ($2x10^6$) or isolated mitochondria or mitochondrial membranes (50 µg) were loaded in the O2k Oxygraph chambers with Mir05 buffer (Oroboros Instruments) and 20 µg digitonin (only for cells); substrates and inhibitors were added sequentially. Glutamate (10 mM) and malate (0.5 mM) were added to activate CI-dependent respiration and rotenone (0.5 µM) was added to suppress it. Succinate (10 mM) was then added to activate CII-dependent respiration, and subsequently 1 mM ADP was added to activate state 3 respiration; both of which were inhibited by adding 2.5 µM antimycin A. Lastly, 0.5 mM N, N, N′, N′-tetramethyl-p-phenylenediamine (TMPD) and 2 mM ascorbate were added to activate CIV-dependent respiration, which was later suppressed by addition of 100 mM sodium azide. In some experiments, 10 µM cyt c was included in the sequence. In some other experiments, 100 nM nigericin or 100 nM monensin was added after succinate.

$Na^+$-containing respiration buffer was prepared by replacing potassium phosphate with sodium phosphate in the Mir05 formula (Oroboros instruments). $Na^+$-free respiration buffer was made with phosphoric acid and buffered to physiological pH with N-methyl D-glucamine (NMDG). Both buffers were compensated in osmolality and ionic strength using NMDG. In addition, in these experiments the acidic forms of substrates, appropriately resuspended in $Na^+$-free respiration solution and buffered to reach respiration solution pH, were used to avoid the introduction of $Na^+$ into the $Na^+$-free respiration buffer or additional $Na^+$ into the $Na^+$-containing buffer. Mitochondrial membrane potential was calibrated according to the manufacturer's instructions.

##### Blue native gel electrophoresis

Complex and supercomplex levels, compositions, and distributions were analyzed in isolated mitochondria by BN-PAGE[28] Mitochondrial proteins were solubilized with 10% digitonin (4 g/g; Sigma-Aldrich D5628) and separated on 3–13%-gradient Blue native gels. Gradient gels (1.5-mm thick) were prepared using a gradient former connected to a peristaltic pump.

##### SDS gel electrophoresis

Mitochondria were resuspended in RIPA buffer (1% Triton-X-100, 50 mM Tris-HCl pH 7.4, 50 mM sodium chloride, 0.5% sodium deoxycholate, and 5 mM EDTA) supplemented with a protease inhibitors cocktail mix (Sigma P8340). Samples were then incubated at 4ºC on a rotating wheel for 15 minutes and centrifuged for 15 minutes at 13,000 x g at 4ºC. Supernatants were collected and transferred to fresh 1.5 ml Eppendorf tubes. Protein concentration was quantified by the Bradford assay.

The levels and distribution of Opa1 were analyzed by sodium dodecyl sulphate (SDS) gel electrophoresis (SDS-PAGE) on 7.5% gels. Loading buffer (50 mM Tris-HCl pH 6.8, 2% SDS, 10% glycerol, 1% 2-mercaptoethanol, 0.02% bromophenol blue) was added to samples, which were incubated for 5 minutes at 95ºC.

##### Immunodetection of single proteins, complexes, and supercomplexes

BN-PAGE or SDS-PAGE proteins were electroblotted using a Mini Trans-Blot Cell system (Bio-Rad) onto polyvinylidene difluoride (PVDF) transfer membrane (Immobilon-FL, 0.45 µm; Merck Millipore, IPFL00010) for 1 hour at 100 V in transfer buffer (48 mM Tris, 39 mM glycine, and

20% ethanol). Non-specific binding sites were blocked by incubating membranes with phosphate-buffered saline (PBS) containing 5% bovine serum albumin (BSA) for 1 hour at RT. For protein detection, membranes were incubated in blocking buffer containing primary antibody overnight at 4ºC. After three washes with PBS 0.1 % Tween-20 (PBS-T) for 30 min, membranes were incubated with appropriate secondary antibodies for 1 hour at RT. Membranes were then washed twice with PBS-T and once with PBS. To study complex and supercomplex assembly, the PVDF membrane was sequentially probed with antibodies against CI (anti-NDUFA9, Abcam ab14713), CIII (anti-UQCR2, Proteintech, 14742-1 AP) and CII (anti-Fp70, Invitrogen, 459200). Parallel BNGE western blots were performed to probe CIV (anti-COI, Invitrogen, MTCO1 459600) and CV (anti-ATPβ, Abcam, Ab 14730). To study Opa1 processing, the PVDF membrane was sequentially probed with antibodies against Opa1 (anti-Opa1, Abcam, ab42364) and Fp70 (anti-Fp70, Invitrogen, 459200). Fiji software was used for Opa1 quantification. Antibody binding was detected by fluorescence as previously described[29].

**Measurement of mitochondrial ATP synthesis**

ATP synthesis was assessed in isolated mitochondria (1.5 to 15 μg mitochondrial protein) using succinate as substrate (plus rotenone) in the presence of ADP in a kinetic luminescence assay based on the luciferin/luciferase reaction[30].

**Measurements of mitochondrial matrix $Na^+$ and $H^+$**

The probe-loading protocol was adapted from[31]. Isolated mitochondria from cells were incubated for 20 min at 37 ºC with 10 μM 2',7'-bis-(2-carboxyethyl)-5-(and-6)-carboxyfluorescein-acetoxymethyl ester (BCECF-AM; for matrix $H^+$) or with 1,3-benzenedicarboxylic acid, 4,4'-[1,4,10-trioxa-7,13-diazacyclopentadecane-7,13-diylbis(5-methoxy-6,12-benzofurandiyl)]bis-,tetrakis [(acetyloxy)methyl] ester (SBFI-AM; for matrix $Na^+$). This was followed by two rounds of centrifugation (12000 g, 4ºC, 5 min) and resuspension in sucrose buffer. The last resuspension was in $Na^+$-free respiration buffer (Extended Data Figure 10a and b).

Probe-loaded mitochondria were distributed in p96 wells in $Na^+$-free buffer, unless otherwise indicated. Measurements were made with a Fluoroskan Ascent microplate reader (Thermo Fisher Scientific) at 390/485 nm excitation pair for BCECF-AM and 355/390 nm excitation pair for SBFI-AM. Emission was recorded at 530 nm emission for BCECF-AM and SBFI-AM. Substrates and inhibitors were subsequently added as in the $O_2$ consumption experiments, without ADP, TMPD, ascorbate, or sodium azide. The $H^+$ pumping rate was obtained from the slope after adding respiratory substrates; the slope was flat after the addition of the corresponding inhibitors. For the measurement of reverse NHE activity using BCECF-AM, 10 mM NaCl was added after succinate. For measurement of forward NHE activity and $Na^+$ gradient using SBFI-AM, 10 mM NaCl was added at the beginning of the experiment.

Calibration was performed by adding equal amounts of mitochondria from the same cell type to several wells containing a graded series of pH or $Na^+$ concentration in the presence of 1 μM nigericin, 1 μM monensin, and 1 μM gramicidin.

**Spectrophotometric swelling assay in isolated mitochondria**

The protocol was adapted to the analysis of cell-culture-derived mitochondria from[32]. Briefly, passive mitochondrial swelling was recorded in 50 μg of mitochondria resuspended in 133 mM sodium acetate (for NHE activity), potassium acetate (for KHE activity) or ammonium aceate (for maximal swelling), 0.2 mM Tris-EGTA, pH 7.0 at room temperature. Absorbance was recorded at 550 nm in a UV/VISJASCO spectrophotometer (Thermo Fisher Scientific).

In some experiments, sodium acetate was exchanged for sodium chloride as a swelling control. In others, before addition of mitochondria to the sodium acetate buffer, mitochondrial CI was activated by incubating the sample with glutamate/malate in Miro05 buffer for 5 min at 37ºC. Alternatively, mitochondrial CI was deactivated by incubating the mitochondria without substrates in Miro05 buffer for 15 min at 37ºC (Extended Data Figure 10c)

**Measurement of $Na^+$ gradient in whole cells by confocal microscopy**

Cytosolic and mitochondrial $Na^+$ were detected as in[29]. Briefly, cybrids were plated the day before experiments, washed three times with Hank's balanced salt solution with $Ca^{2+}$/$Mg^{2+}$/glucose (HBSS + Ca/Mg + glucose), and incubated with 5 µM Asante NaTRIUM Green-2-Acteoxymethyl ester (ANG2-AM) for 30 min at 37 °C in the dark. ANG2-AM was washed out, and new HBSS + Ca/Mg + glucose was added, including 1 µM CoroNa Red. Cells were further incubated for 30 min at 37 °C in the dark. After this, the medium was changed again, cells were washed once with HBSS + Ca/Mg + glucose, and the plate was placed on the automated stage of a Leica SP-5 confocal microscope for live imaging. The planes were focused for image capture, and images were taken with a ×63 objective. Experiments started and ended at 20% $O_2$ and 5% $CO_2$. Loaded cells were excited with an argon/krypton laser using the 496-nm line for ANG2-AM and the 514-nm line for CoroNa Red. Fluorescence emission of ANG2-AM was detected in the 515–550-nm range, whereas the emission of CoroNa Red was detected in the 555–575-nm range.

*In situ* calibration of the same cells was performed after a two-wash step with $Na^+$-free HBSS + Ca/Mg + glucose. Increasing $Na^+$ concentrations, starting from 0 mM $Na^+$, were then applied in the presence of 1 µM nigericin, 1 µM monensin, and 1 µM gramicidin, and images were taken with the excitation/emission wavelengths indicated above. Calibration solutions were equilibrated for at least 5 min.

**Measurement of mitochondrial $Ca^{2+}$ and pH, and cytosolic $Ca^{2+}$ in whole cells by confocal microscopy**

The protocol followed was exactly as in[29]. Briefly, cells were transiently transfected with Cepia2mt (for mitochondrial $Ca^{2+}$), mitosypHer (for mitochondrial pH) or cytoGECO (for cytosolic $Ca^{2+}$) and were all imaged under a Leica SP5 microscope. Cellular and mitochondrial $Ca^{2+}$ entry was stimulated by addition of Histamine 100 µM and calibrated using ionomycin 2 µM and EDTA 10 mM. pH calibration was performed with HBSS + Ca/Mg + glucose + monensin 1 µM, nigericin 1 µM and gramicidin 1 µM at different pHs.

**Measurement of $\Delta\Psi$mt in whole cells by confocal microscopy**

Cells were seeded the day prior experimentation. Then, cells were incubated with TMRM 50 nM, for non-quenching mode, for 30 min before imaging in a Leica SP5 microscope. Samples were excited with an argon/krypton laser using the 543-nm line and emission was detected in the 555–595-nm range. Calibration was performed following guidelines and indications published elsewhere[33].

**Measurement of mitochondrial volume**

For the determination of mitochondrial volume 50 µg of mitochondria and 1 mL Mir05 respiration buffer were placed in a cuvette, and the absorbance at 550 nm was recorded with a UV/VISJASCO spectrophotometer (Thermo Fisher Scientific). Parallel measurements were taken at steady state and in mitochondria respiring from different substrates for 5 min at 37 ºC. A valinomycin technical control was included to determine whether this technique was suitable for the measurement of mitochondrial volume changes, with positive results.

**Transmission electron microscopy**

Cells were analysed by electron microscopy as previously described[29]. Thin sections including mitochondria were imaged with a JEM1010 electron microscope (Jeol).

**Proteomic analysis**

Protein extracts of whole cells or isolated mitochondria from WT, ND6$^{KD}$, ND4$^{KO}$, NDUFS4$^{WT}$, NDUFS4$^{KO}$, hWT, LHON or pure CI samples were obtained by homogenization with ceramic beads (MagNa Lyser Green Beads, Roche, Germany) in CS buffer (Pipes pH6.8, MgCl2, NaCl, EDTA, sucrose, SDS, sodium orthovanadate; Biochain Institute, Inc. #K3013010-5) freshly supplemented with protease and phosphatase inhibitors. Extracted proteins (around 200 µg) were subjected to in-filter reduction and alkylation using iodoacetamide followed by trypsin digestion (Nanosep Centrifugal Devices with Omega Membrane-10K, PALL), and the resulting peptides were TMT-labeled according to the manufacturer's instructions. Labeled peptides were loaded and washed on Evotips for chromatographic separation by an evosep one HPLC system (30 SPD method, with Endurance Column 15 cm x 150 µm ID, 1.9 µm beads-EV1106, Evosep). Mass spectra were acquired in a data-dependent manner, with an automatic switch between MS and MS/MS using a top-speed method and dynamic exclusion. MS spectra were collected in the Orbitrap analyzer using a mass range of 375–1500 m/z at 60,000 resolution. HCD fragmentation was performed at 33 eV of normalized collision energy and MS/MS spectra were analyzed at 30,000 resolution in the Orbitrap.

Proteins were identified with the SEQUEST HT algorithm integrated in Proteome Discoverer 2.5 (Thermo Scientific). MS/MS scans were searched against a pig reference target-decoy protein database (human_pig_202105_pro-sw-tr.target-decoy.fasta), (296316 sequences in total). For database searching, parameters were selected as follows: trypsin digestion with 2 maximum missed cleavage sites, precursor mass tolerance of 2 Da, and a fragment mass tolerance of 0.03 Da. Methionine oxidation (+15.994915 Da) and asparagine and glutamine deamidation (+0.984016 Da) were set as variable modifications, whereas cysteine carbamidomethylation (+57.021464 Da) and TMT labeling (+229.162932 Da) at peptide N-terminal ends and Lys residues were considered fixed modifications. False discovery rates (FDR) for peptide identifications were calculated by the refined method[34][35] after additional filtering for a precursor mass tolerance of 10 ppm[35][36]. A 1% FDR was used as criterion for peptide identification.

Quantitative information from TMT reporter intensities was integrated from the spectrum level to the peptide level and then to the protein level based on the WSPP model[36][37] [37][38], using the GIA integration algorithm[38][39].

To note, taking into account that the protein extraction protocol is not specific for membrane proteins, the peptides of these proteins are poorly represented in this study.. This includes ND6 or ND4 in all models analyzed.

**Purification of Porcine Heart Complex I**

Intact mitochondria were isolated from fresh homogenized fresh pig heart (Icarlopsa Slaughterhouse in Tarancón, Spain) in 0.25 M sucrose, 10 mM Tris-HCl (pH 7.4) and washed twice in the same medium before fragmentation as previously described[39]. The pellet was resuspended in 50 mM TrisHCl (pH 8.0), 20% (v/v) glycerol, frozen in liquid nitrogen, and stored at -80ºC. The Complex I was purified using an adapted multiple column strategy as described by Letts et al.[40] alternating anion exchange with buffer A (20 mM Tris-HCl, pH 7.4, 10% (v/v) glycerol, 1 mM EDTA, 1 mM DTT, 0.1 mg/ml DOPC, and 0.1 % DDM) and buffer B(20 mM Tris-HCl, pH 7.4, 10% (v/v) glycerol, 1 mM EDTA, 1 mM DTT, 0.1 mg/ml DOPC, and 0.1 % DDM, 1M NaCl) followed by size exclusion chromatography (SEC) using the SEC buffer (20 mM HEPES, pH 7.4, 2 mM EDTA, 10% glycerol, 50 mM NaCl, 0.1 mg/ml DOPC/CL (4:1) and 0.05% DDM).

Briefly, to solubilize complex I from the mitochondrial membranes DDM (10% w/v) was added to dropwise to a final concentration of 1% DDM and incubated for 30 minutes on an orbital shaker at 4 °C. To remove the non-solubilized material samples were centrifuged at 80,000 × g for 20 min (Beckman MLA80 rotor, Optima-MAX XP ultracentrifuge).

Next, the obtained Supernatant was filtered (0.45 µm pore size) and loaded onto 20-ml Mono Q HR anion-exchange column (GE Healthcare) previously equilibrated with equilibration buffer (20 mM Tris-HCl, pH 7.4, 10% (v/v) glycerol, 1 mM EDTA, 1 mM DTT, 0.1 mg/ml DOPC, and 0.1 % DDM, 50 mM NaCl). After the application of the mitochondrial extract, the Mono Q column was washed with 11 ml of 5% buffer B, followed with a 11-ml linear gradient of 5 - 23% buffer B, and finally with 68 ml of 23% buffer B. Complex I was then eluted with a 89-ml linear gradient of 23–30% buffer B followed with 40 ml of 100% buffer B to elute any protein from the column. The Q-Sepharose gradient was run at 1.0 ml/min at 4 °C. Complex I containing fractions were pooled based on NADH/FeCy activity and concentrated with centriprep-10 K Centrifugal Filters (Milipore) to a final volume of 10 ml. This sample was loaded onto a Hiprep 26/60 Sephacryl S 300 HR column (GE Healthcare) equilibrated with SEC buffer and eluted overnight at 0.35 ml/min at 4 °C.

Complex I containing fractions were identified by spectrophotometrically CI activity and BN-PAGE, followed by CI in-gel activity.

#### Reconstitution of Complex I into proteoliposomes

The fraction chosen for reconstitution was the corresponding to the CI in-gel activity BN-PAGE band which showed more CI abundancy and a specific activity band at the level of CI (fraction 10). This fraction was also analyzed by mass spectrometry to confirm CI purity. Purified Complex I was reconstituted into proteoliposomes using the rapid detergent dilution method[41]. For this purpose, 100 µl of solubilized Complex I (0.2 mg/mL) was mixed with 20 µl of synthetic DOPC/CL (4:1 w/w; 20 mg/mL) and incubated for 30 min on ice. Subsequently, the mixture was diluted by the addition of 4 ml of reconstitution buffer (50 mM Tris-HCl, pH 7,4) and left on ice for additional 20 min. Complex I-containing proteoliposomes (C1-PL) were collected by high-speed centrifugation (30 min; 400000 g; 4°C) and resuspended in 100 µl (initial volume of the purified Complex I) of reconstitution buffer. The reconstituted C1-PL were aliquoted and used for further experiments.

#### Measurement of Pure Complex I NHE activity into proteoliposomes

CI-containing proteoliposomes were either reconstituted with acridine orange (AO; 10 µM) or only buffer. Proteoliposomes with AO were incubated for 5 min in reconstitution buffer, measurement started once the plate reached 37ºC and NaCl 30 mM was added during the measurement. Measurements were made with a Fluoroskan Ascent microplate reader (Thermo Fisher Scientific) at 495/525 nm excitation/emission filter pair. Unloaded proteoliposomes were added to the preheated reconstitution solution containing 30 mM NaCl and either 10 µM of SBFI-AM or 10 µM BCECF-AM during the fluorometric measurement (Extended Data Figure 10d). Measurements were made with a Fluoroskan Ascent microplate reader (Thermo Fisher Scientific) at 390/485 nm excitation pair for BCECF-AM and 355/390 nm excitation pair for SBFI-AM. Emission was recorded at 530 nm emission for BCECF-AM and SBFI-AM.

**Analysis of mitochondrial morphology by confocal microscopy**

Glass cover slips were consecutively washed with 100% EtOH and PBS 1x and irradiated with UV light for 20 min to sterilize them. Once dry, one cover slip was placed per well on a P12 cell culture plate. 30,000 cells were plated per well in complete DMEM and were left incubating at 37ºC overnight. The following day, cells were incubated for 20 min at 37ºC in DMEM containing 33 nM Mitotracker® Deep Red FM, after which the medium was discarded and exchanged for fresh DMEM in which the cells remained for 2 h at 37ºC. Finally, cells were fixed in 4% PFA for 15 min at 37ºC, followed by three washes in PBS 1x. Cells were stained with DAPI (1:1000) and mounted using ProLong® Gold Antifade Reagent (Invitrogen). Confocal fluorescence microscopy images were obtained at a Leica SP5 inverted confocal microscope using HCX PL APO lambda blue 63x 1.40 oil objectives and a digital zoom of 2x. Z-stack images (step size: 0.35 µm) were acquired with LAS-AF 2.6.0 software (Leica Microsystems).

Image analysis was performed in FIJI 1.53t using the Mitochondria Analyzer plugin (PMID: 31846372). Maximum projection images of Mitotracker signal were thresholded with the 2D Batch Analysis function with the following parameters: subtract background (rolling: 2.0), sigma filter plus (2.0 radius), enhance local contrast (max slope: 3.0), weighted mean threshold (block size: 1.0, C-value: 5.0), despeckle and remove outliers (pixels: 4.0). Individual cells were isolated from whole thresholded images, and size parameter of mitochondria were calculated on a per cell basis.

**Measurement of $H^+$ movement by electrophysiology**

Measurements were made in an O2k Oxygraph instrument (Oroboros Instruments) with the attached pH electrode module. Isolated intact mitochondria (100 µg), liposomes (100 µl) or CI-containing proteoliposomes (100 µl) were loaded in the O2k Oxygraph chambers with a low-buffering capacity buffer (2 mM imidazole, 140.66 mM KCl 140.66 mM, 0.49 $MgCl_2$ and 0.1 mM EDTA, pH 7.1 (adjusted at 25º C). pH electrode was calibrated in the day of the experiment and the buffer pH recorded before and after the addition of 30 mM NaCl.

**Statistical analysis**

Statistical analyses and graphics were produced with GraphPad Prism 8 software. Datasets were compared by t test, analysis of variance (ANOVA), or nonparametric analysis as appropriate and with P values adjusted for multiple tests. P values from each comparison are shown in every graph. All results are presented as mean ± SD or mean ± SEM

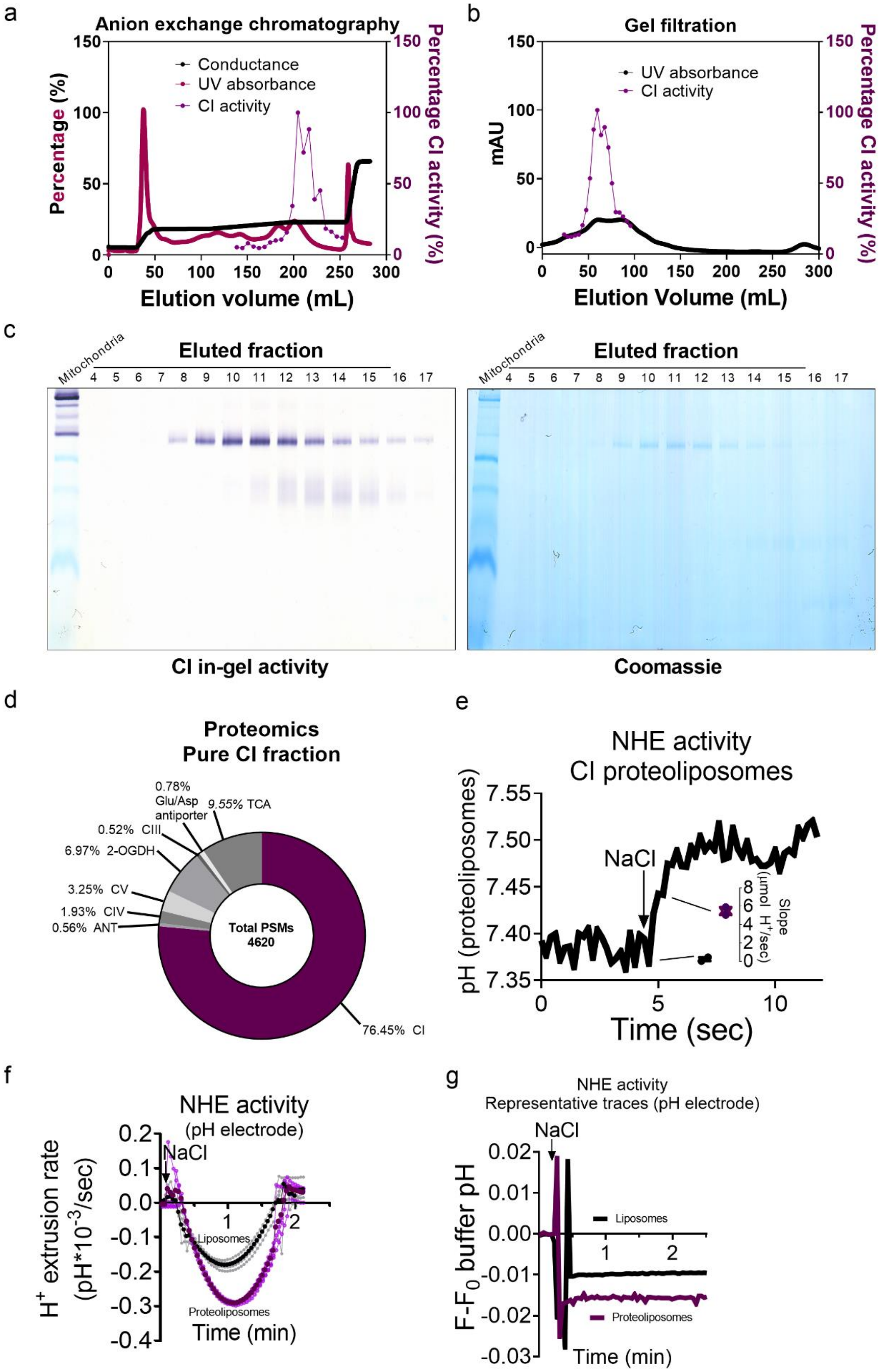
a
Anion exchange chromatography
Percentage (%)
Percentage CI activity (%)
Conductance
UV absorbance
CI activity
Elution volume (mL)
b
Gel filtration
mAU
Percentage CI activity (%)
UV absorbance
CI activity
Elution Volume (mL)
c
Mitochondria
Eluted fraction
4 5 6 7 8 9 10 11 12 13 14 15 16 17
CI in-gel activity
Coomassie
d
Proteomics
Pure CI fraction
0.78% Glu/Asp antiporter
9.55% TCA
0.52% CIII
6.97% 2-OGDH
3.25% CV
1.93% CIV
0.56% ANT
Total PSMs 4620
76.45% CI
e
NHE activity
CI proteoliposomes
pH (proteoliposomes)
NaCl
Slope (µmol H+/sec)
Time (sec)
f
NHE activity
(pH electrode)
H+ extrusion rate (pH*10-3/sec)
NaCl
Liposomes
Proteoliposomes
Time (min)
g
NHE activity
Representative traces (pH electrode)
NaCl
Liposomes
Proteoliposomes
F-F0 buffer pH
Time (min)

**Extended Data Figure 1. Pure CI reconstituted into proteoliposomes is a $Na^+/H^+$ antiporter.** (**a**) Anion exchange chromatography profile showing the percentage of conductance, UV absorbance and CI activity. (**b**) Gel filtration profile showing the UV absorbance and percentage of CI activity of the samples previously enriched by anion exchange chromatography. (**c**) BN-PAGE CI in-gel activity (left) and Coomassie staining (right) of the eluted fractions from the gel filtration containing CI. Isolated pig heart mitochondria were loaded as a control to corroborate sole migration of CI in the eluted fractions. (**d**) Proteomic analysis of the fraction used for proteoliposome reconstitution. (**e**) Fluorescence of CI-containing proteoliposomes loaded with 10 µM BCECF was measured before and after the addition of NaCl 10 mM. Inset shows the quantification of two independent experiments. (**f**) Buffer pH was measured with a pH electrode in a solution containing CI-containing proteoliposomes or liposomes before and after the addition of NaCl 30 mM (n=4). (**g**) Representative raw traces of buffer pH from (**f**).

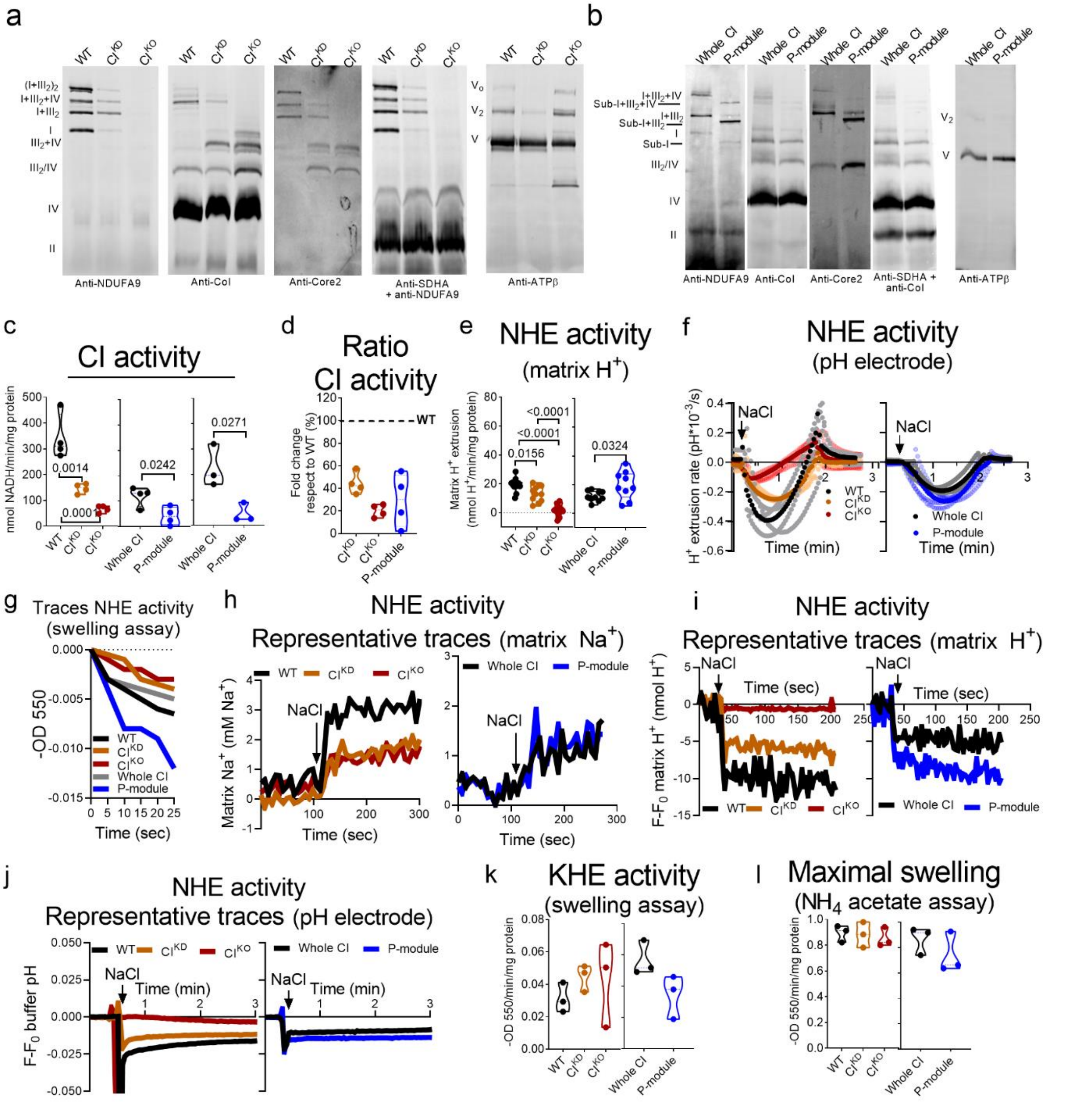


**Extended Data Figure 2. Differential CI subunit suppression translates into either loss or gain mNHE function**. (**a**) Blue-native electrophoresis (BN-PAGE) of mitochondria from WT, $CI^{KD}$, and $CI^{KO}$ cybrid cell lines (n=3 per genotype) immunoblotted to detect subunits of CI (Anti-NDUFA9), CIV (Anti-CoI), CIII (Anti-Core2), CII (Anti-SDHA), and CV (Anti-ATPβ). * marks an unspecific band. (**b**) BN-PAGE of mitochondria from $NDUFS4^{WT}$ (Whole CI) and $NDUFS4^{KO}$ (P-module) MAFs (n=2) blotted against the same epitopes as in (a). The lateral labels in (a) and (b) indicate the migrated positions of all detected complexes and supercomplexes. (**c**) Rotenone-sensitive NADH-decylCoQ oxidoreductase activity in WT, $CI^{KD}$ and $CI^{KO}$ mitochondrial membranes (n=4). (**d**) Ratio of the CI activity of $CI^{KD}$, $CI^{KO}$ and $NDUFS4^{KO}$ (P-module) with respect their isogenic control (n=4). (**e**) Mitochondrial matrix $H^+$ extrusion measured with BCECF-

AM in WT, $CI^{KD}$, $CI^{KO}$, $NDUFS4^{WT}$ (Whole CI) and $NDUFS4^{KO}$ (P-module) intact mitochondria respiring on succinate and 1 µM rotenone (n=11; n=10 in Whole CI and n=9 in P-module). (**f**) Buffer pH was measured with a pH electrode in a solution containing 100 µg of WT, $CI^{KD}$, $CI^{KO}$, $NDUFS4^{WT}$ (Whole CI) and $NDUFS4^{KO}$ (P-module) intact mitochondria before and after the addition of NaCl 30 mM. (**g**) Representative traces of passive NHE activity swelling assay of WT, $CI^{KD}$, $CI^{KO}$, $NDUFS4^{WT}$ and $NDUFS4^{KO}$ intact mitochondria. (**h**) Representative traces of mitochondrial matrix $Na^+$ measured with SBFI-AM in WT, $CI^{KD}$, $CI^{KO}$, $NDUFS4^{WT}$ (Whole CI) and $NDUFS4^{KO}$ (P-module) intact mitochondria before and after the addition of 10 mM NaCl. (**i**) Representative traces of mitochondrial matrix $H^+$ measured with BCECF-AM in WT, $CI^{KD}$, $CI^{KO}$, $NDUFS4^{WT}$ and $NDUFS4^{KO}$ intact mitochondria respiring on succinate and 1 µM rotenone before and after the addition of 10 mM NaCl. (**j**) Representative raw traces of buffer pH from (**f**). Passive KHE activity swelling assay in WT, $CI^{KD}$, $CI^{KO}$, $NDUFS4^{WT}$ and $NDUFS4^{KO}$ intact mitochondria (n=3). (**k**) Maximal swelling assay in WT, $CI^{KD}$, $CI^{KO}$, $NDUFS4^{WT}$ (Whole CI) and $NDUFS4^{KO}$ (P-module)intact mitochondria was performed in $NH_4$ acetate buffer (n=3).

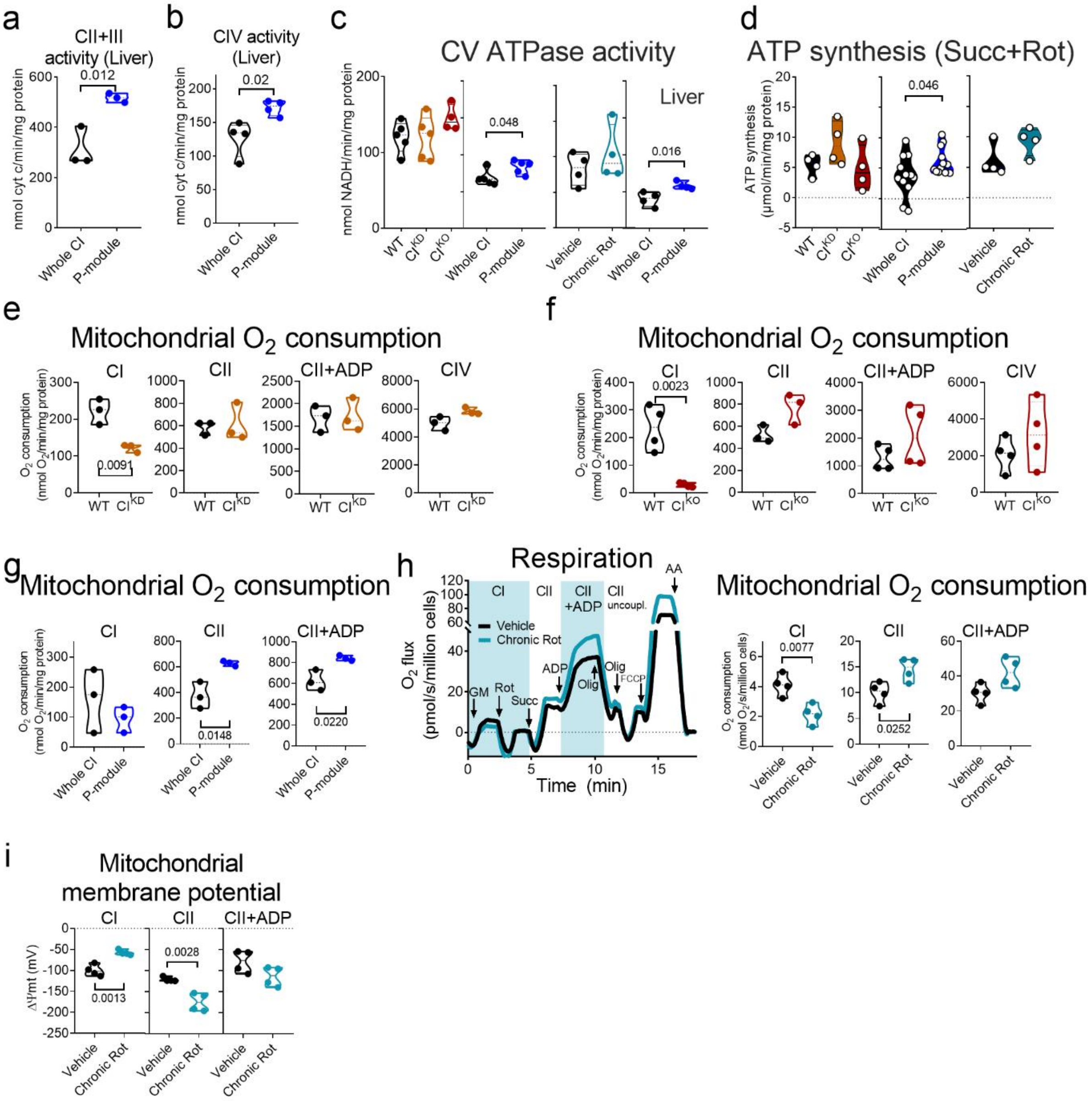


**Extended Data Figure 3. Bioenergetics analysis of isolated mitochondria from all CI deficiency models.** (**a**) Antimycin A (AA)-sensitive succinate-cyt c oxidoreductase activity in NDUFS4$^{WT}$ (Whole CI) and NDUFS4$^{KO}$ (P-module) mouse liver mitochondrial membranes (n=3). (**b**) KCN-sensitive cyt c oxidase activity in NDUFS4$^{WT}$ (Whole CI) and NDUFS4$^{KO}$ (P-module) mouse liver mitochondrial membranes (n=3). (**c**) Oligomycin (Olig)-sensitive ATPase activity in WT (n=5), CI$^{KD}$ (n=5), CI$^{KO}$ (n=4), Vehicle-treated (n=4), Chronic rotenone-treated (n=4) and NDUFS4$^{WT}$ (n=5) and NDUFS4$^{KO}$ (n=5) mitochondrial membranes from MAFs and Liver. (**d**) Olig-sensitive ATP synthase activity in WT, CI$^{KD}$, CI$^{KO}$, Vehicle-treated, Chronic rotenone-treated (n=4) and NDUFS4$^{WT}$ (Whole CI) and NDUFS4$^{KO}$ (P-module) (n=12) isolated mitochondria in the presence of 1 µM Rot and respiring on succinate. (**e-g**) Quantification of respiratory rates of WT and CI$^{KD}$ (**e**), WT and CI$^{KO}$ (**f**) and NDUFS4$^{WT}$ (Whole CI) and NDUFS4$^{KO}$ (P-module) (n=3) (**g**) isolated mitochondria respiring on CI substrates (CI inset), CII substrates (CII inset), CII substrates +ADP (CII+ADP inset), and CIV substrates (CIV inset) (n=3). (**h**) (left) Oxygen consumption rates were measured in permeabilized WT cells that had been treated with vehicle or Chronic Rot (n=4). Insets and acronyms are the same as those in Fig. 2 and sections (e-g); FCCP, carbonyl cyanide-p-trifluoromethoxyphenylhydrazone (n=4). (**i**) Calibrated TMRM signal in permeabilized WT cells that had been treated with vehicle or Chronic Rot, respiring on CI substrates (CI inset), CII substrates (CII inset), or CII substrates +ADP (CII+ADP inset).

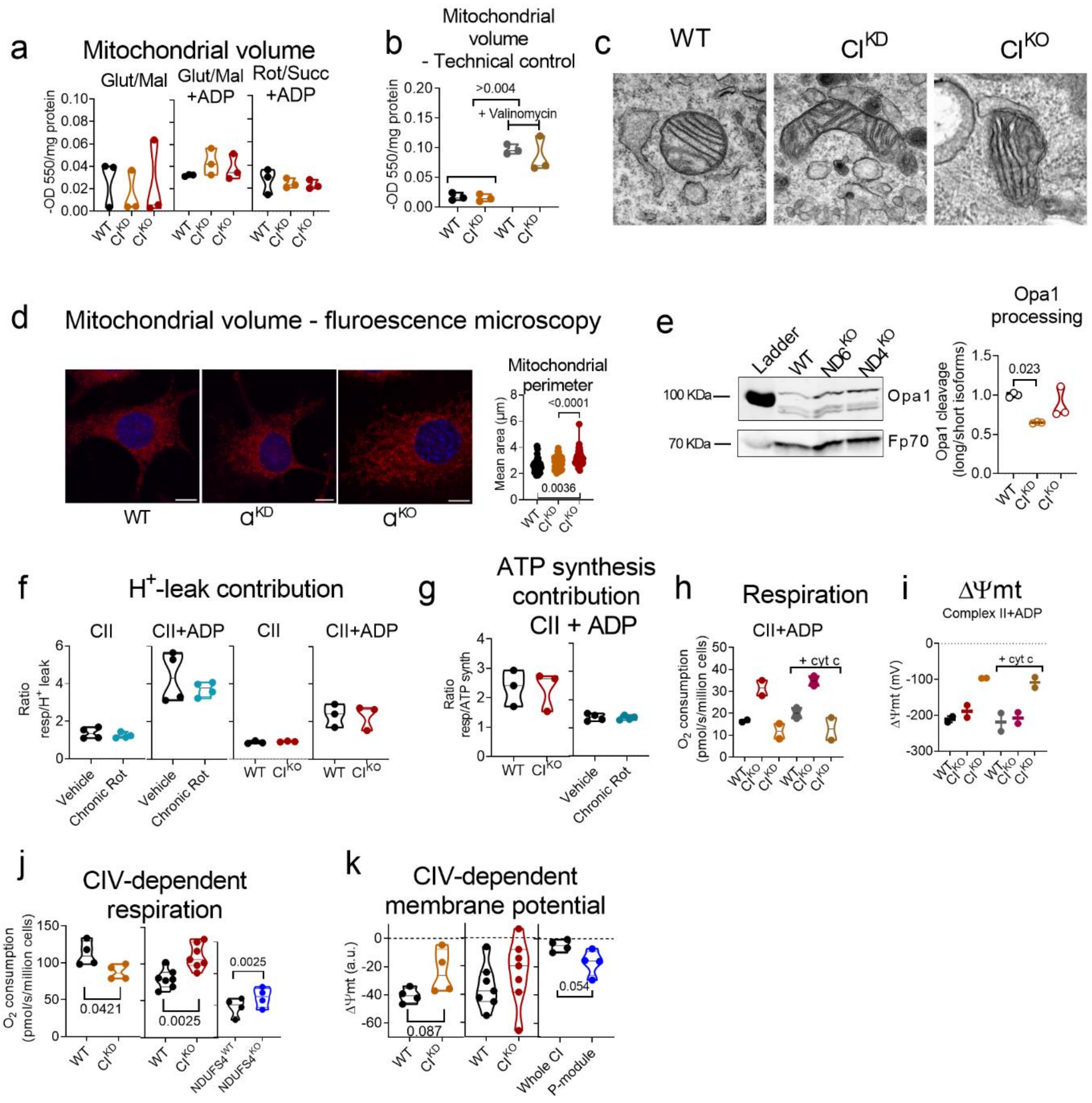


**Extended Data Figure 4. CI$^{KD}$ and CI$^{KO}$ mitochondria have similar size and membrane permeability than WT and maintain their bioenergetic footprint under CIV substrates.** (**a**) Mitochondrial volume measured by absorbance at 550 nm in the presence of CI substrates (Glut/Mal), CI substrates + ADP (Glut/Mal+ADP), or CII substrates + 1 µM rotenone (Rot/Succ+ADP) in WT, CI$^{KD}$ and CI$^{KO}$ mitochondria. (**b**) WT and CI$^{KD}$ mitochondrial volume in the absence or presence of valinomycin. (**c**) Transmission electron microscopy (TEM) imaging of mitochondria in WT, CI$^{KD}$, and CI$^{KO}$ cells (n=2). (**d**) Mitotracker Deep-Red and DAPI staining of WT, CI$^{KD}$ and CI$^{KO}$ were analyzed by confocal microscopy. Inset: mitochondrial area was calculated with a Mitochondria Analyzer plugin (n=37). (**e**) Western blot analysis of OPA1 expression in WT, CI$^{KD}$, and CI$^{KO}$ isolated mitochondrial fractions; anti-Fp70 was used as a loading control. Inset: shows quantification of three independent experiments. (**f**) $H^+$ leak contribution to baseline CII respiration +/- ADP in Vehicle and Chronic Rot-treated, as well as in WT and $CI^{KO}$ permeabilized cells, calculated as the ratio of baseline CII respiration (without ADP) to $H^+$ leak. (**g**) ATP synthesis contribution to CII respiration in Vehicle and Chronic Rot-treated, as well as in WT and $CI^{KO}$ permeabilized cells, calculated as the ratio of coupled CII respiration (with ADP) to ATP synthesis. (**h**) Oxygen consumption rates in WT, $CI^{KD}$, and $CI^{KO}$ permeabilized cells respiring on Succ + Rot + ADP in the absence or presence of cyt c. (**i**) Calibrated TMRM signal in WT, $CI^{KD}$, and $CI^{KO}$ permeabilized cells respiring on Succ + Rot + ADP in the absence or presence of cyt c. (**j**) Quantification of respiratory rates of WT vs $CI^{KD}$ (n=4), WT vs $CI^{KO}$ (n=7) and $NDUFS4^{WT}$ (Whole CI) and $NDUFS4^{KO}$ (P-module) (n=4) permeabilized cells respiring on CIV substrates. (**k**) Calibrated TMRM signal in WT vs $CI^{KD}$ (n=4), WT vs $CI^{KO}$ (n=7) and $NDUFS4^{WT}$ vs $NDUFS4^{KO}$ (n=4) permeabilized cells respiring on CIV substrates.

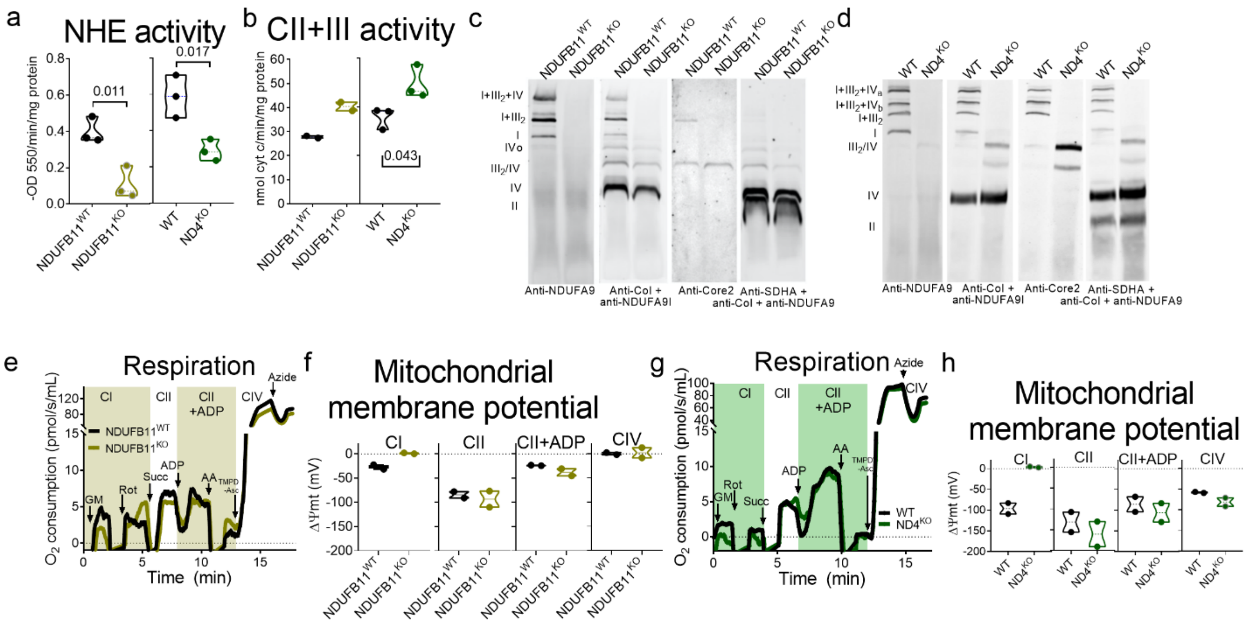


**Extended Data Figure 5. Deletion of a nuclear encoded CI P-module subunit resembles the bioenergetic footprint of a P-module mtDNA mutant**. (**a**) Passive NHE activity swelling assay in NDUFB11$^{WT}$, NDUFB11$^{KO}$, WT and ND4$^{KO}$ intact mitochondria (n=3). (**b**) Antimycin A (AA)-sensitive succinate-cyt c oxidoreductase activity in NDUFB11$^{WT}$, NDUFB11$^{KO}$, WT and ND4$^{KO}$ mitochondrial membranes (n=2 in NDUFB11$^{KO}$ and n=3 in ND4$^{KO}$). (**c-d**) BN-PAGE of mitochondria from NDUFB11$^{WT}$, NDUFB11$^{KO}$ (**c**), WT and ND4$^{KO}$ (**d**; n=2) blotted against the same epitopes as in Extended Data Fig 2a, except for CV. The lateral labels indicate the migrated positions of all detected complexes and supercomplexes. (**e**) Oxygen consumption rates were measured in NDUFB11$^{WT}$ and NDUFB11$^{KO}$ isolated mitochondria (n=2). Insets and acronyms are the same as those in Figure 2. (**f**) Calibrated TMRM signal in NDUFB11$^{WT}$ and NDUFB11$^{KO}$ isolated mitochondria (n=2), respiring on CI substrates (CI inset), CII substrates (CII inset), or CII substrates +ADP (CII+ADP inset). (**g**) Oxygen consumption rates were measured in WT and ND4$^{KO}$ isolated mitochondria (n=2). Insets and acronyms are the same as those in Figure 2. (**f**) Calibrated TMRM signal in in WT and ND4$^{KO}$ isolated mitochondria (n=2), respiring on CI substrates (CI inset), CII substrates (CII inset), or CII substrates +ADP (CII+ADP inset).

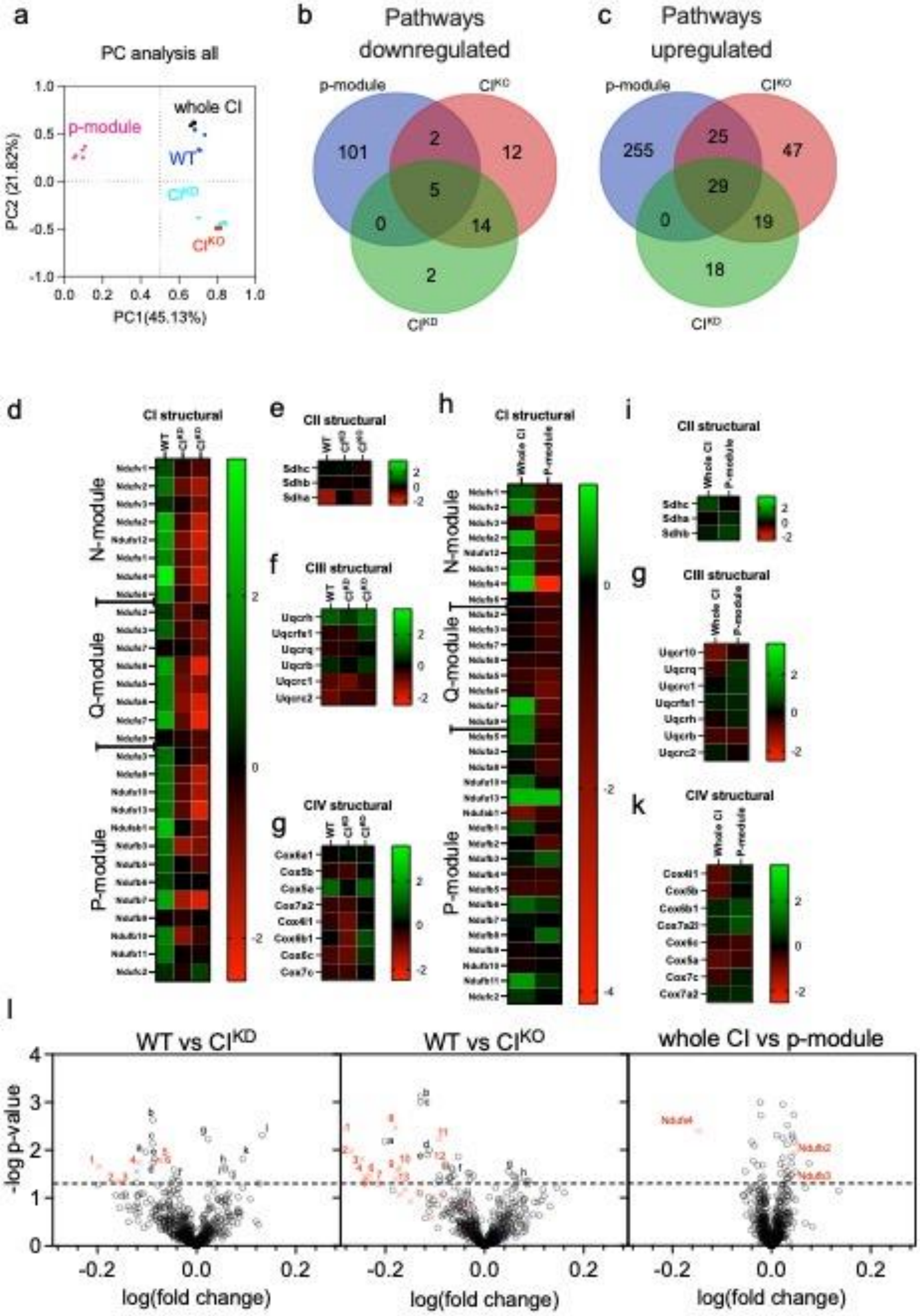
a
PC analysis all
p-module
whole CI
WT*
CI^KD
CI^KO
PC2 (21.82%)
PC1(45.13%)
b
Pathways downregulated
p-module
CI^KO
CI^KD
101
2
12
5
0
14
2
c
Pathways upregulated
p-module
CI^KO
CI^KD
255
25
47
29
0
19
18
d
CI structural
N-module
Q-module
P-module
e
CII structural
f
CIII structural
g
CIV structural
h
CI structural
N-module
Q-module
P-module
i
CII structural
g
CIII structural
k
CIV structural
l
WT vs CI^KD
WT vs CI^KO
whole CI vs p-module
-log p-value
log(fold change)
Ndufa4
Ndufb2
Ndufb3

**Extended Data Figure 6. Proteomics analysis of WT, $CI^{KD}$, $CI^{KO}$, $NDUFS4^{WT}$ (Whole CI) and $NDUFS4^{KO}$ (P-module) isolated mitochondria. (a)** Principal component analysis using the whole quantified protein dataset. **(b-c)** Venn diagrams comparing the Gene Ontology pathways significantly downregulated **(b)** or upregulated **(c)** estimated by Gene Set Enrichment Analysis (GSEA). (**d-k**) Mean relative protein abundances of (**d**) CI subunits in WT, $CI^{KD}$, and $CI^{KO}$ mitochondria; (**e**) CII subunits in WT, $CI^{KO}$, and $CI^{KO}$ mitochondria; (**f**) CIII subunits in WT, $CI^{KD}$, and $CI^{KO}$ mitochondria; (**g**) CIV subunits in WT, $CI^{KD}$, and $CI^{KO}$ mitochondria; (**h**) of CI subunits in $NDUFS4^{WT}$ and $NDUFS4^{KO}$ mitochondria; (**i**) of CII subunits in $NDUFS4^{WT}$ (Whole CI) and $NDUFS4^{KO}$ (P-module)mitochondria; (**j**) of CIII subunits in $NDUFS4^{WT}$ (Whole CI) and $NDUFS4^{KO}$ (P-module)mitochondria and (**k**) of CIV subunits in $NDUFS4^{WT}$ (Whole CI) and $NDUFS4^{KO}$ (P-module)mitochondria. (**l**) Volcano plot analysis showing relative abundance changes of all mitochondrial proteins detected in WT vs $CI^{KD}$; in WT vs $CI^{KO}$; in $NDUFS4^{WT}$ (Whole CI) and $NDUFS4^{KO}$ (P-module). n=3 for all experiments. Statistical significance of changes between groups were evaluated by Student's t-test. Labels in WT vs $CI^{KD}$: (1) NDUFS4, (2) NDUFA7, (3) NDUFB3, (4) NDUFS6, (5) NDUFS2, (6) NDUFB11; (blue cross): NDUFAF8; (a) Timm44, (b) Fkbp10, (c) Acadm, (d) Dbi, (e) Dcxr, (group f) Acp6, Dbt, Mrpl33, D2hgdh, (g) Trap1, (h) Acad10, (i) Pitrm1, (j) Smim20, (k) Aldhl2 and (l) Acad1. Labels in WT vs $CI^{KO}$: (1) NDUFS4, (2) NDUFA7, (3) NDUFS8, (4) NDUFB7, (5) NDUFA12, (6) NDUFA2, (7) NDUFA13, (8) NDUFS6, (9) NDUFA8, (10) NDUFA10, (11) NDUFB5, (12) NDUFB11, (13) NDUFAB1; (blue cross): NDUFAF8; (a) Acot2, (b) Triap1, (c) Cisd3, (d) Mrpl12, (e) Acadm, (group f) Aldh9a1, Fxn, Acadl, Spire1, Fdps, Dld, Rdh13, Rot1, Tom34, Htaip2, Acap1, (g) Mul1, (group h) Gatd3a, Mmab, Rmdn3, Acadsb and Maoa. Labels in Whole vs P-module: (1) NDUFS4, (2) NDUFB2, (3) NDUFB3; Downregulated: Lylpa1, Agpa5, Prkaca, Clpp, Dld, Nsun2, Slc25a12, CoQ3, Mfn2; Upregulated: Dmac1, Slc25a15, Bcs1l, Ptrh1, Mrpl34, Auh, Timm13, Fastk, Timm22, Sdhb, Timm29, C2hgdh, Mrpl10, Mrps27, Slc25a11, Timm8a1, Dmac2, Hsd17b10, Tstd3 and Chchd4.

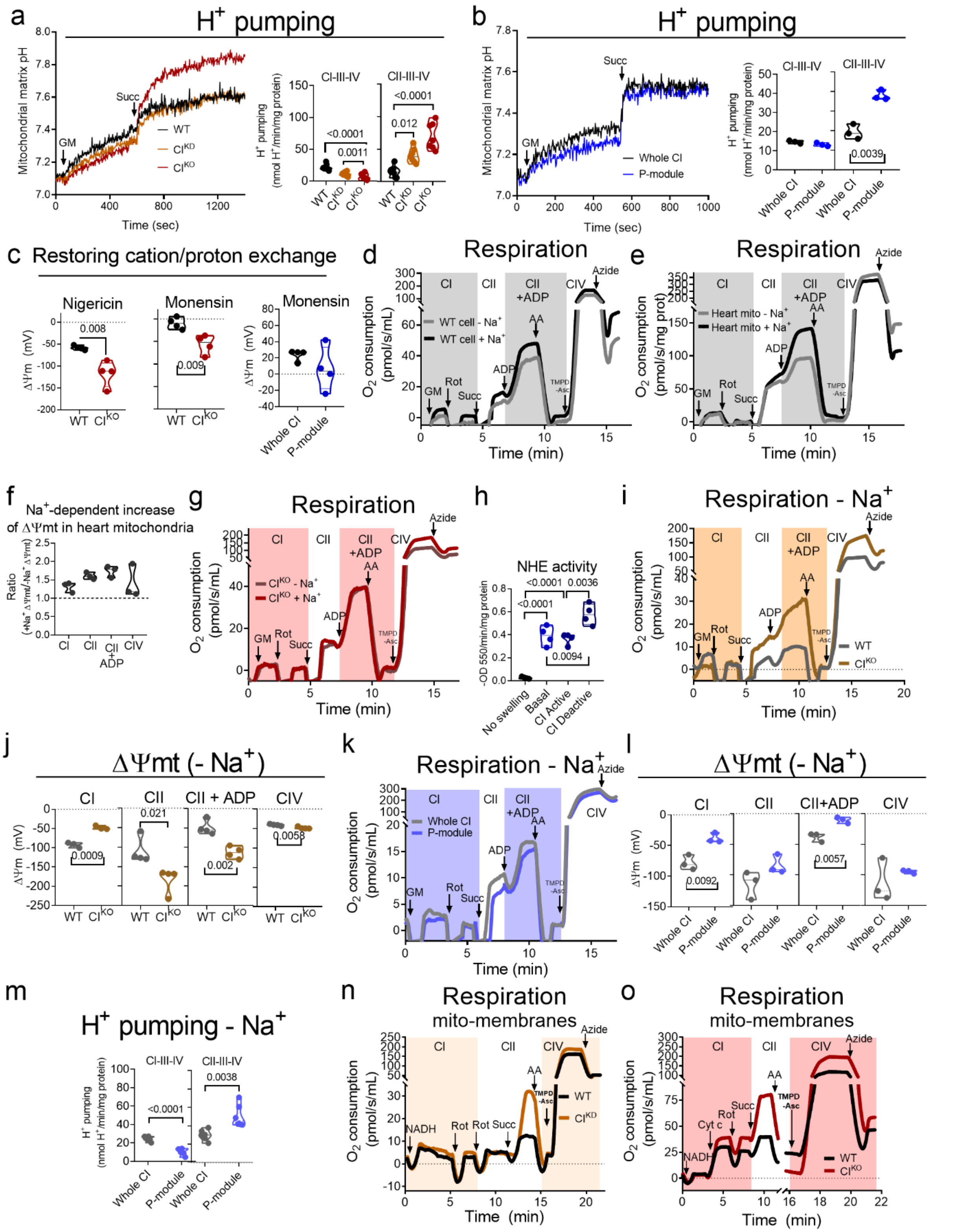


**Extended Data Figure 7. CI-NHE activity-dependent Na$^+$ gradient contributes to total ΔΨmt.** (**a**) Calibrated BCECF-AM signals of WT, CI$^{KD}$ and CI$^{KO}$ isolated mitochondria under CI (Inset

CI-CIII-CIV) and CII substrates (Inset CII-CIII-CIV). (**b**) Calibrated BCECF-AM signals of NDUFS4$^{WT}$ (Whole CI) and NDUFS4$^{KO}$ (P-module) isolated mitochondria under CI (Inset CI-CIII-CIV) and CII substrates (Inset CII-CIII-CIV). (**c**) Subtraction of calibrated TMRM signal of WT and CI$^{KO}$ mitochondria after and before the addition of nigericin (100 nM) or monensin (100 nM) and only monensin (100 nM) in NDUFS4$^{WT}$ (Whole CI) and NDUFS4$^{KO}$ (P-module) isolated mitochondria respiring in Rot/Succ + ADP. (**d** and **e**) Oxygen consumption rates measured in WT (**d**; n=4) and C57BL/6N heart (**e**; n=3) mitochondria in $Na^+$-containing buffer (black) and $Na^+$-free buffer (gray; n=4). (**f**) Relative contribution of $Na^+$ to $\Delta\Psi$mt in C57BL/6N heart mitochondria respiring on CI, CII, CII+ADP, or CIV substrates, calculated as the ratio of the calibrated TMRM signal in $Na^+$-containing to that in $Na^+$-free buffer. (**g**) Oxygen consumption rates measured in CI$^{KO}$ mitochondria in $Na^+$-containing buffer (red) and $Na^+$-free buffer (brown; n=4). (**h**) Mouse liver mitochondrial NHE activity in different CI activation states (n=4). (**i**) Oxygen consumption rates measured in WT and CI$^{KO}$ mitochondria in $Na^+$-free buffer (n=4). (**j**) Calibrated TMRM signal in WT and CI$^{KO}$ mitochondria respiring on CI substrates (CI inset), CII substrates (CII inset), CII substrates +ADP (CII+ADP inset), and CIV substrates (CIV inset) in a $Na^+$-free buffer. (**k**) Oxygen consumption rates measured in NDUFS4$^{WT}$ (Whole CI) and NDUFS4$^{KO}$ (P-module) mitochondria in $Na^+$-free buffer (n=3). (**l**) Calibrated TMRM signal in NDUFS4$^{WT}$ (Whole CI) and NDUFS4$^{KO}$ (P-module) mitochondria respiring on CI substrates (CI inset), CII substrates (CII inset), CII substrates +ADP (CII+ADP inset), and CIV substrates (CIV inset) in a $Na^+$-free buffer (n=3). (**m**) Calibrated BCECF-AM signals of NDUFS4$^{WT}$ (Whole CI) and NDUFS4$^{KO}$ (P-module) isolated mitochondria under CI (Inset CI-CIII-CIV) and CII substrates (Inset CII-CIII-CIV) in a $Na^+$-free buffer (n=3). (**n**) Oxygen consumption rates in WT and CI$^{KD}$ frozen-thawed-permeabilized mitochondria in the presence of cyt c (n=3). (**o**) Oxygen consumption rates in WT and CI$^{KO}$ frozen-thawed-permeabilized mitochondria in the presence of cyt c (n=3). Labels are the same as in Fig. 2.

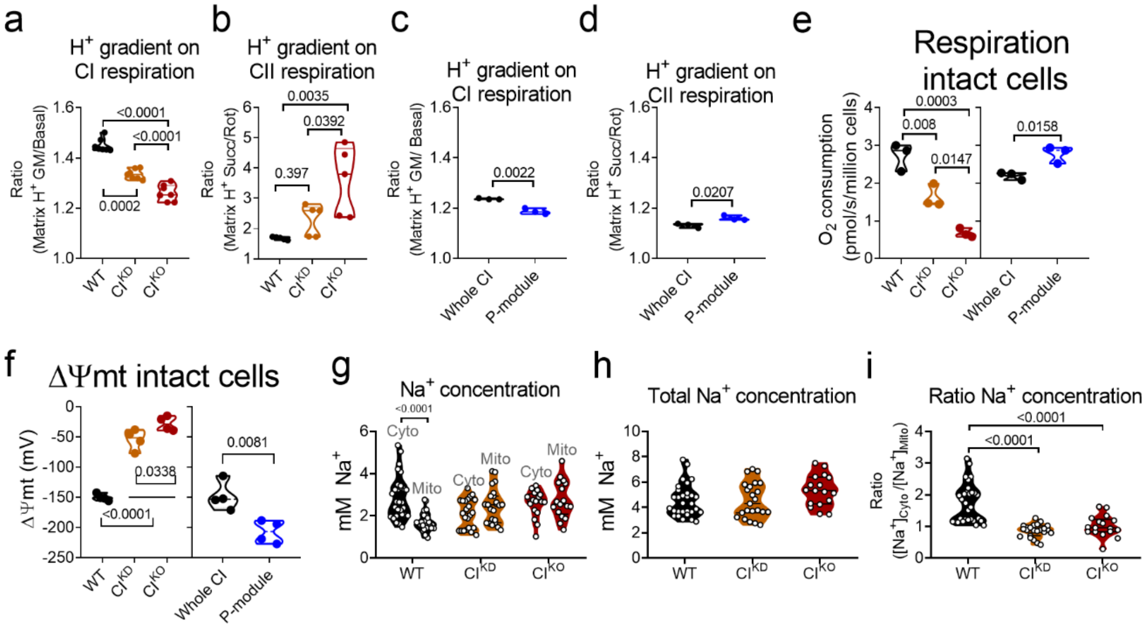


**Extended Data Figure 8. The mitochondrial Na$^+$ gradient parallels the H$^+$ gradient and contributes to respiratory rates and ΔΨmt in intact cells.** (**a** and **b**) H$^+$ gradient in WT, CI$^{KD}$ and CI$^{KO}$ isolated mitochondria incubated with BCECF-AM and respiring on GM (**a**) or succinate + rotenone 1μM (**b**), calculated as the ratio of matrix H$^+$ concentrations determined after and before addition of rotenone (**a**) or AA (**b**). (**f**) H$^+$ gradient in NDUFS4$^{WT}$ (Whole CI) and NDUFS4$^{KO}$ (P-module) isolated mitochondria incubated with BCECF-AM and respiring on GM, calculated as the ratio of matrix H$^+$ concentrations determined after and before addition of rotenone. (**c** and **d**) H$^+$ gradient in NDUFS4$^{WT}$ (Whole CI) and NDUFS4$^{KO}$ (P-module) isolated mitochondria incubated with BCECF-AM and respiring on GM (**c**) or succinate + rotenone 1μM (**d**), calculated as the ratio of matrix H$^+$ concentrations determined after and before addition of rotenone (**c**) or AA (**d**). (**e**) Oxygen consumption in WT, CI$^{KD}$, CI$^{KO}$, NDUFS4$^{WT}$ (Whole CI) and NDUFS4$^{KO}$ (P-module) intact cells. (**f**) Calibrated TMRM signal of WT, CI$^{KD}$, CI$^{KO}$, NDUFS4$^{WT}$ (Whole CI) and NDUFS4$^{KO}$ (P-module) intact cells imaged by confocal microscopy. (**g**) Cytosolic and mitochondrial Na$^+$ concentrations in WT, CI$^{KD}$, and CI$^{KO}$ intact cells measured by confocal microscopy. (**h**) Total Na$^+$ concentrations calculated as the sum of cytosolic and mitochondrial Na$^+$ concentrations, measured by confocal microscopy. (**i**) Mitochondrial Na$^+$ gradient calculated as the ratio between cytosolic and mitochondrial Na$^+$ concentrations measured by confocal microscopy.

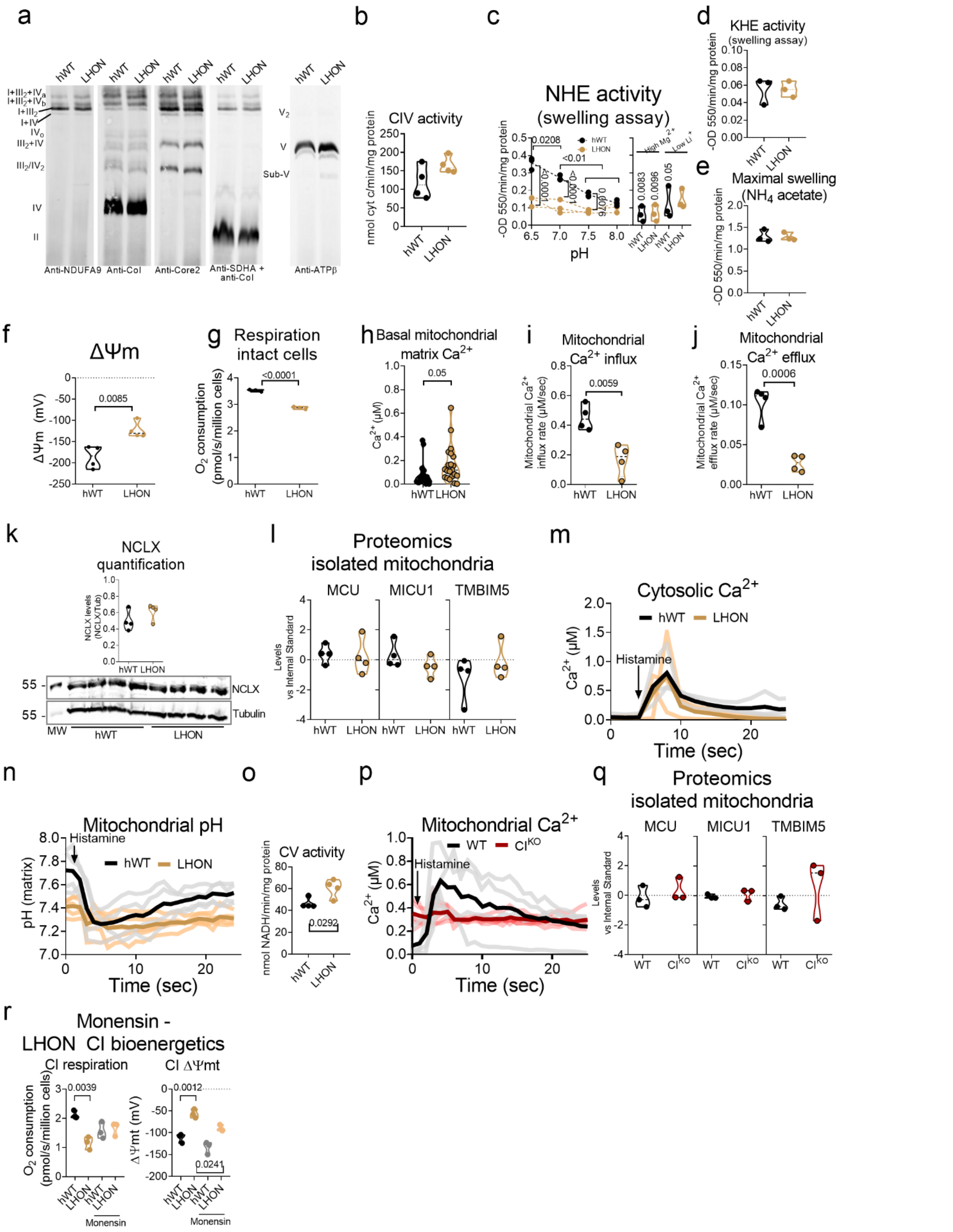
a
hWT LHON hWT LHON hWT LHON hWT LHON hWT LHON
I+III2+IVa
I+III2+IVb
I+III2
I+IV
IVo
III2+IV
III2/IV2
IV
II
V2
V
Sub-V
Anti-NDUFA9
Anti-CoI
Anti-Core2
Anti-SDHA + anti-CoI
Anti-ATPβ
b
CIV activity
nmol cyt c/min/mg protein
hWT LHON
c
NHE activity
(swelling assay)
-OD 550/min/mg protein
0.0208
<0.01
<0.0001
hWT
LHON
High Mg2+
Low Li+
0.0083
0.0096
0.05
pH
d
KHE activity
(swelling assay)
-OD 550/min/mg protein
hWT LHON
e
Maximal swelling
(NH4 acetate)
-OD 550/min/mg protein
hWT LHON
f
ΔΨm
ΔΨm (mV)
0.0085
hWT LHON
g
Respiration
intact cells
O2 consumption
(pmol/s/million cells)
<0.0001
hWT LHON
h
Basal mitochondrial
matrix Ca2+
Ca2+ (μM)
0.05
hWT LHON
i
Mitochondrial
Ca2+ influx
Mitochondrial Ca2+
influx rate (μM/sec)
0.0059
hWT LHON
j
Mitochondrial
Ca2+ efflux
Mitochondrial Ca2+
efflux rate (μM/sec)
0.0006
hWT LHON
k
NCLX
quantification
NCLX levels
(NCLX/Tub)
hWT LHON
55
NCLX
55
Tubulin
MW
hWT
LHON
l
Proteomics
isolated mitochondria
MCU
MICU1
TMBIM5
Levels
vs Internal Standard
hWT LHON hWT LHON hWT LHON
m
Cytosolic Ca2+
hWT
LHON
Histamine
Ca2+ (μM)
Time (sec)
n
Mitochondrial pH
Histamine
hWT
LHON
pH (matrix)
Time (sec)
o
CV activity
nmol NADH/min/mg protein
0.0292
hWT LHON
p
Mitochondrial Ca2+
WT
CIKO
Histamine
Ca2+ (μM)
Time (sec)
q
Proteomics
isolated mitochondria
MCU
MICU1
TMBIM5
Levels
vs Internal Standard
WT CIKO WT CIKO WT CIKO
r
Monensin -
LHON CI bioenergetics
CI respiration
CI ΔΨmt
O2 consumption
(pmol/s/million cells)
0.0039
ΔΨmt (mV)
0.0012
0.0241
hWT LHON hWT LHON
Monensin

**Extended Data Figure 9. The specific defect of CI-NHE in LHON-causing m.11778 causes defects in $Ca^{2+}$ management.** (**a**) Blue-native electrophoresis (BN-PAGE) of mitochondria from hWTand LHON cybrid cell lines (n=2) immunoblotted to detect subunits of CI (Anti-NDUFA9), CIV (Anti-CoI), CIII (Anti-Core2), CII (Anti-SDHA), and CV (Anti-ATPβ). (**b**) KCN-sensitive cyt c oxidation in mitochondrial membranes from hWT and LHON mitochondria (n=4). (**c**) Passive swelling NHE activity in hWT and LHON was assessed in different buffer pHs and in the presence of 50 mM $MgCl_2$ or with 2 mM LiCl (n=3). (**d**) Passive KHE activity swelling assay in hWT and LHON intact mitochondria (n=3). (**e**) Maximal swelling assay in hWT and LHON intact mitochondria was performed in $NH_4$ acetate buffer (n=3). (**f**) Calibrated TMRM signal of hWT and LHON intact cells imaged by confocal microscopy (n=4). (**g**) Oxygen consumption in hWT and LHON intact cells (n=3). (**h**). Basal mitochondrial $Ca^{2+}$ levels were measured by confocal microscopy in hWT and LHON cells transfected with Cepia2mt (n=18 in hWT and n=21 in LHON). (**i**) Mitochondrial $Ca^{2+}$ influx was calculated from the signals in Fig. 4k (n=4). (**j**) Mitochondrial $Ca^{2+}$ efflux was calculated from the signals in Fig. 4k (n=4). (**k**) SDS-PAGE western blot showing the levels of NCLX and Tubulin from whole hWT and LHON cell extracts (MW: Molecular weight). Inset: Ratio NCLX/Tubulin of four independent biological replicates. (**l**) Proteomic analysis showing the relative abundances of MCU, MICU2 and TMBIM5 relative to internal standard in hWT and LHON isolated mitochondria. (**m**) Cytosolic $Ca^{2+}$ levels measured by confocal microscopy of hWT and LHON cells transfected with cytoGECO before and after the addition of histamine. (**n**) Mitochondrial pH levels measured by confocal microscopy of hWT and LHON cells transfected with mito-sypHer before and after the addition of histamine. (**o**) Oligomycin (Olig)-sensitive ATPase activity in hWT and LHON mitochondrial membranes. (**p**) Mitochondrial $Ca^{2+}$ was measured with calibrated Cepia2mt in WT vs $CI^{KO}$ cells before and after stimulation with 100 µM histamine (n=4 in WT and n=5 in $CI^{KO}$). (**q**) Proteomic analysis showing the relative abundances of MCU, MICU and TMBIM5 relative to internal standard in WT and $CI^{KO}$ isolated mitochondria. (**r**) CI-dependent respiratory rates (left) and $\Delta\Psi$mt (right) isolated mitochondria treated with Vehicle or 50 nM monensin.

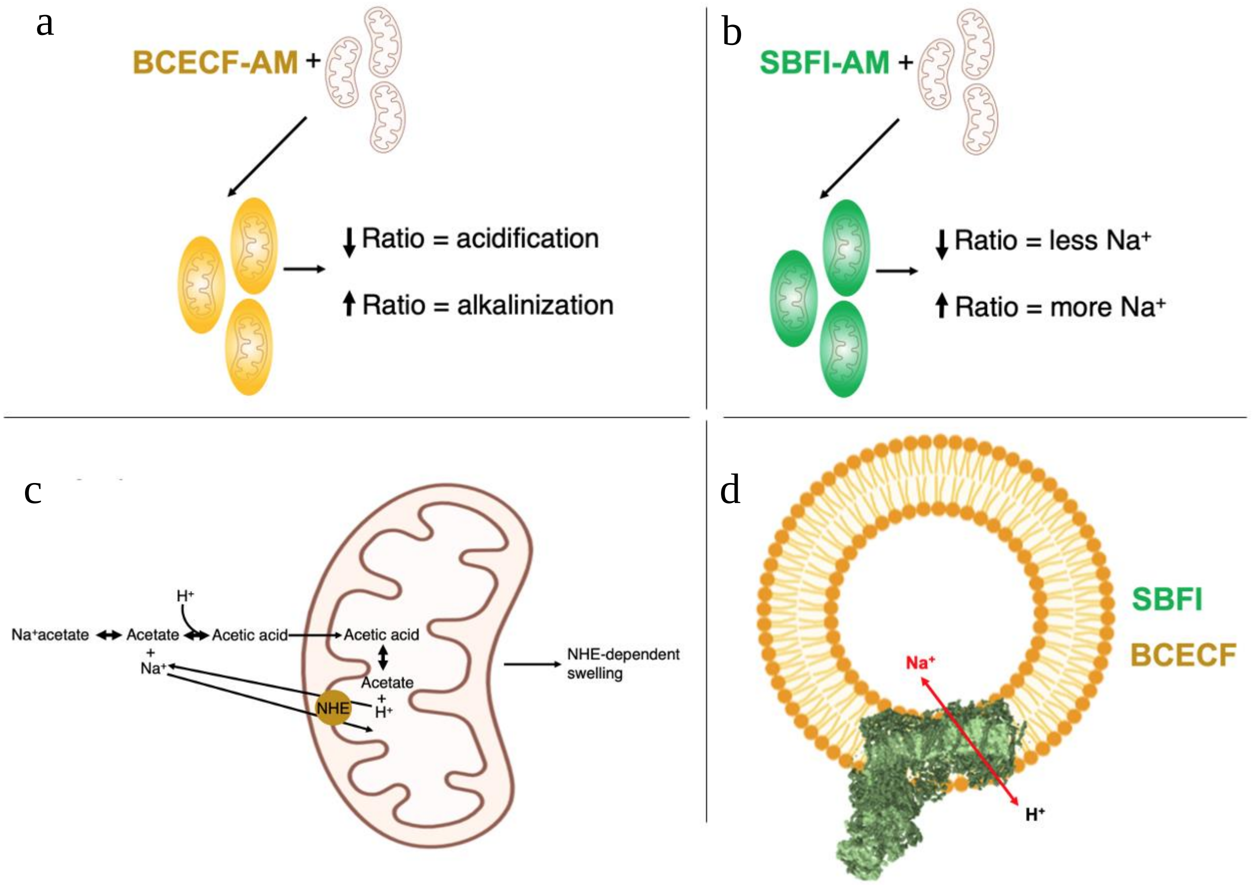


**Extended Data Figure 10**. **Schemes showing the methodological or physical basis of some protocols used in this study.** **(a)** BCECF-AM or **(b)** SBFI-AM were incubated 20 min with isolated mitochondria and, once the probe was loaded, mitochondrial suspension was washed twice by centrifugation. **(c)** Isolated mitochondria are treated with a buffer containing high amounts of $Na^{+}$-acetate. Acetate dissociates from $Na^{+}$ and some becomes protonated due buffer equilibrium. Acetic acid then crosses the mitochondrial membranes and becomes again deprotonated due to the buffer's equilibrium. The released $H^{+}$ in the matrix and the Na+ in the buffer are used by the mNHE. Then, mitochondrial $Na^{+}$ entry promotes swelling, which is a direct readout of mNHE. **(d)** CI-reconstituted proteoliposomes were added to a physiological solution containing $Na^{+}$, SBFI and BCECF. Then, changes in $Na^{+}$ amount and $H^{+}$ were monitored by fluorescence.

**Extended Data Table 1. Summary of bioenergetics characterization of CI-deficiency models.**

| Comparisons vs Control 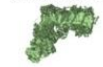 | **CII+III activity** | **CIV activity** | **CV activity** | **Respiration (CII- and CIV-depedent)** | **$\Delta\Psi$mt** | **mNHE** | **Succinate-driven $H^+$ pumping** |
|---|---|---|---|---|---|---|---|
| **$CI^{KD}$** 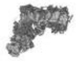 | Higher (Fig. 2a) | Equal (Fig. 2b) | Equal (ED Fig. 3c-d) | Equal (Fig. 2c and ED Fig. 3e) | Equal (Fig. 2d) | Lower (Fig. 1b and d and ED Fig 2) | Higher (ED Fig. 7a) |
| **$CI^{KO}$** | Higher (Fig. 2a) | Higher (Fig. 2b) | Equal (ED Fig. 3c-d) | Equal (Fig. 2e and ED Fig. 3f) | Equal (Fig. 2f) | Lower (Fig. 1b and d and ED Fig 2) | Higher (ED Fig. 7a) |
| **P-module** 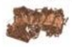 | Higher (Fig. 2a) | Higher (Fig. 2b) | Higher (ED Fig. 3c-d) | Higher (Fig. 2g and ED Fig. 3g) | Hyperpolarized (Fig. 2h) | Higher (Fig. 1b and d and ED Fig 2) | Higher (ED Fig. 7b) |
| **Chronic Rot** 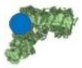 | Higher (Fig. 2a) | Equal (Fig. 2b) | Equal (ED Fig. 3c-d) | Higher (ED Fig. 3h) | Hyperpolarized (ED Fig. 3i) | - | - |

**Extended Data Table 2. Summary of total and relative contributions of the $Na^+$ gradient to ΔΨmt in the presence of different respiratory substrates and oxidizing conditions.**

| | Estimation approach | $Na^+$-dependent ΔΨmt (mV) | Contribution to ΔΨmt (%) |
|---|---|---|---|
| **CI respiration** (Isolated mitochondria) | ΔΨmt measurement in WT± $Na^+$ | -92.78 ± 27.31 | 34.72 ± 6.35 |
| **CII respiration** (Isolated mitochondria) | ΔΨmt measurement in WT± $Na^+$ | -157.46 ± 37.89 | 34.49 ± 5.74 |
| **CII+ ADP respiration** (Isolated mitochondria) | TMRM fluorescence ± monensin | -86.14 ± 16.97 | 53.11 ± 5.79 |
| | ΔΨmt measurement in WT± $Na^+$ | -111.06 ± 16.93 | |


26 Fernandez-Vizarra, E. *et al.* Isolation of mitochondria for biogenetical studies: An update. *Mitochondrion* **10**, 253-262 (2010). https://doi.org:10.1016/j.mito.2009.12.148

27 Morava, E. *et al.* Clinical and biochemical characteristics in patients with a high mutant load of the mitochondrial T8993G/C mutations. *Am J Med Genet A* **140**, 863-868 (2006). https://doi.org:10.1002/ajmg.a.31194

28 Wittig, I., Braun, H. P. & Schagger, H. Blue native PAGE. *Nat Protoc* **1**, 418-428 (2006). https://doi.org:10.1038/nprot.2006.62

29 Hernansanz-Agustin, P. *et al.* Na(+) controls hypoxic signalling by the mitochondrial respiratory chain. *Nature* **586**, 287-291 (2020). https://doi.org:10.1038/s41586-020-2551-y

30 Vives-Bauza, C., Yang, L. & Manfredi, G. Assay of mitochondrial ATP synthesis in animal cells and tissues. *Methods Cell Biol* **80**, 155-171 (2007). https://doi.org:10.1016/S0091-679X(06)80007-5

31 Dlaskova, A., Hlavata, L., Jezek, J. & Jezek, P. Mitochondrial Complex I superoxide production is attenuated by uncoupling. *Int J Biochem Cell Biol* **40**, 2098-2109 (2008). https://doi.org:10.1016/j.biocel.2008.02.007

32 Douglas, M. G. & Cockrell, R. S. Mitochondrial cation-hydrogen ion exchange. Sodium selective transport by mitochondria and submitochondrial particles. *J Biol Chem* **249**, 5464-5471 (1974).

33 Gerencser, A. A. *et al.* Quantitative measurement of mitochondrial membrane potential in cultured cells: calcium-induced de- and hyperpolarization of neuronal mitochondria. *J Physiol* **590**, 2845-2871 (2012). https://doi.org:10.1113/jphysiol.2012.228387

34 Navarro, P. & Vazquez, J. A refined method to calculate false discovery rates for peptide identification using decoy databases. *J Proteome Res* **8**, 1792-1796 (2009). https://doi.org:10.1021/pr800362h

35 Bonzon-Kulichenko, E., Garcia-Marques, F., Trevisan-Herraz, M. & Vazquez, J. Revisiting peptide identification by high-accuracy mass spectrometry: problems associated with the use of narrow mass precursor windows. *J Proteome Res* **14**, 700-710 (2015). https://doi.org:10.1021/pr5007284

36 Trevisan-Herraz, M. *et al.* SanXoT: a modular and versatile package for the quantitative analysis of high-throughput proteomics experiments. *Bioinformatics* **35**, 1594-1596 (2019). https://doi.org:10.1093/bioinformatics/bty815
37 Prieto, G. & Vazquez, J. Protein Probability Model for High-Throughput Protein Identification by Mass Spectrometry-Based Proteomics. *J Proteome Res* **19**, 1285-1297 (2020). https://doi.org:10.1021/acs.jproteome.9b00819
38 Garcia-Marques, F. *et al.* A Novel Systems-Biology Algorithm for the Analysis of Coordinated Protein Responses Using Quantitative Proteomics. *Mol Cell Proteomics* **15**, 1740-1760 (2016). https://doi.org:10.1074/mcp.M115.055905
39 Smith, A. L. [13] Preparation, properties, and conditions for assay of mitochondria: Slaughterhouse material, small-scale. *Methods in Enzymology* **10**, 81-86 (1967). https://doi.org:doi.org/10.1016/0076-6879(67)10016-5
40 Letts, J. A., Degliesposti, G., Fiedorczuk, K., Skehel, M. & Sazanov, L. A. Purification of Ovine Respiratory Complex I Results in a Highly Active and Stable Preparation. *J Biol Chem* **291**, 24657-24675 (2016). https://doi.org:10.1074/jbc.M116.735142
41 Brundage, L., Hendrick, J. P., Schiebel, E., Driessen, A. J. & Wickner, W. The purified E. coli integral membrane protein SecY/E is sufficient for reconstitution of SecA-dependent precursor protein translocation. *Cell* **62**, 649-657 (1990). https://doi.org:10.1016/0092-8674(90)90111-q
